# A graded neonatal mouse model of necrotizing enterocolitis demonstrates that mild enterocolitis is sufficient to activate microglia and increase cerebral cytokine expression

**DOI:** 10.1101/2023.08.03.551849

**Authors:** Cuilee Sha, Trevor Van Brunt, Jacob Kudria, Donna Schmidt, Alisa Yurovsky, Jela Bandovic, Michael Giarrizzo, Joyce Lin, Styliani-Anna Tsirka, Agnieszka B Bialkowska, Lonnie Wollmuth, Esther Speer, Helen Hsieh

## Abstract

**Background:** Necrotizing enterocolitis (NEC) is an inflammatory gastrointestinal process that afflicts approximately 10% of preterm infants born in the United States each year, with a mortality rate of 30%. NEC severity is graded using Bell’s classification system, from stage I mild NEC to stage III severe NEC. Over half of NEC survivors present with neurodevelopmental impairment during adolescence, a long-term complication that is poorly understood but can occur even after mild NEC. Although multiple animal models exist, none allow the experimenter to control nor represent the gradient of symptom severities seen in NEC patients. We bridge this knowledge gap by developing a graded murine model of NEC and studying its relationship with neuroinflammation across a range of NEC severities.

**Methods:** Postnatal day 3 (P3) C57BL/6 mice were fed a formula containing different concentrations (0% control, 0.25%, 1%, 2%, and 3%) of dextran sodium sulfate (DSS). P3 mice were fed every 3 hours for 72-hours. We collected data on weight gain and behavior (activity, response, body color) during feeding. At the end of the experiment, we collected tissues (intestine, liver, plasma, brain) for immunohistochemistry, immunofluorescence, and cytokine and chemokine analysis.

**Results:** Throughout NEC induction, mice fed higher concentrations of DSS died sooner, lost weight faster, and became sick or lethargic earlier. Intestinal characteristics (dilation, color, friability) were worse in mice fed with higher DSS concentrations. Histology revealed small intestinal disarray among mice fed all DSS concentrations, while higher DSS concentrations resulted in reduced small intestinal cellular proliferation and increased hepatic and systemic inflammation. In the brain, IL-2, G-CSF, and CXCL1 concentrations increased with higher DSS concentrations. Although the number of neurons and microglia in the CA1 hippocampal region did not differ, microglial branching was significantly reduced in DSS-fed mice.

**Conclusion:** We characterize a novel graded model of NEC that recapitulates the full range of NEC severities. We show that mild NEC is sufficient to initiate neuroinflammation and microglia activation. This model will facilitate studies on the neurodevelopmental effects of NEC.

## Introduction

Necrotizing enterocolitis (NEC) is the most frequent intestinal emergency among preterm newborns and accounts for ∼125,000 cases per year in North America [1]. Most (70%) NEC patients can be treated medically (bowel rest, antibiotics, and parenteral nutrition), but severely affected patients require surgery for necrotic or perforated bowel [2]. Using Bell’s staging criteria, NEC severity is graded into three stages (mild, moderate, and severe) based on clinical parameters like abdominal distension, hemodynamic stability, radiological findings, and the necessity of surgical intervention [2]. Overall mortality for NEC ranges from 15 to 30%, whereas half of surgical NEC patients die [3–5]. In addition to gastrointestinal (GI) dysfunction, the major long-term complications of NEC are diminished growth and neurodevelopmental impairments, which are found in nearly 60% of NEC survivors [6].

The neurological outcomes following NEC range from mild developmental delays to severe cognitive dysfunction and neuromotor deficits. NEC patients have a 50% increased risk for neurodevelopmental impairments compared to gestational age-matched cohorts of premature patients [6]. These neurological symptoms are associated with changes in cross-sectional imaging that include white matter lesions, cortical thinning, and loss of gray matter volume [7]. Even patients who have had mild NEC experience behavioral problems and learning disabilities in school at a higher rate than premature patients without NEC [8–10].

The pathways underlying the complex relationship between GI pathology and central nervous system (CNS) changes remain incompletely understood. Previous literature has demonstrated changes in brain volume, neuronal cell numbers, inflammatory markers, and microglia activation as a direct consequence of severe NEC in animal models and human postmortem samples [7, 11–14]. However, current models are limited in their ability to study the initial steps of NEC pathogenesis and cannot prospectively induce mild and severe NEC in animal models. A postmortem examination is typically required to correlate NEC colitis severity with central nervous system changes. An animal model that can prospectively control NEC severity would be a valuable tool to study the effects of NEC severities on subsequent neurodevelopmental outcome.

Building upon a NEC model described by Ginzel et al. in 2017 [15], we developed and characterized a graded model of NEC that allows us to control the severity of induced colitis. By adding increasing concentrations of dextran sodium sulfate (DSS), an osmotic agent, to enteral gavage feedings of neonatal mice starting on the third day of life, we demonstrated a graded response in mortality and clinical characteristics, as well as in intestinal morphology and disease appearance. We chose an experimental time period in mouse neuronal development of postnatal days 3 to 6 that correspond with the third trimester of human brain development, thus allowing us to model how NEC affects brain development in preterm infants [16]. In the periphery, DSS supplementation correlated with changes in the concentrations of a wide array of inflammatory indexes in liver and plasma. In the brain, only a small subset of these markers was significantly increased. This novel titratable neonatal NEC model will advance our understanding of the initial stages of NEC, their transition to more advanced NEC stages, and the interplay between the GI, immune, and nervous systems.

## Materials and Methods

### Animals

All animal experiments were performed according to a protocol approved by the Institutional Animal Care and Use Committee (IACUC) of Stony Brook University, Stony Brook, NY, in concordance with the guidelines established by the National Institutes of Health (NIH). Mice (C57BL/6, Charles River, Wilmington, MA) were bred and housed within Stony Brook University’s animal care and research facilities, which are part of the Division of Laboratory Animal Resources (DLAR).

### Necrotizing Enterocolitis (NEC) Induction with Dextran Sodium Sulfate (DSS)

Enterocolitis was induced in mice, as described by Ginzel et al. (2017) [15]. Mice were separated from their mothers and placed in an Ohmeda Medical Ohio® Care Plus Incubator (36.8°C) starting on postnatal day 3 (P3). Animals from the same litter were randomly assigned, without consideration of sex, to control or experimental groups. Two hours after separation from the dam, pups were fed by orogastric gavage every three hours for 72 hours total. Pups were initially fed with 50 μL of Esbilac formula (PetAg, Hampshire, IL) with or without dextran sodium sulfate (DSS, Sigma-Aldrich, St Louis, MO) supplementation. Gavage volumes were increased by 10 μL daily to account for weight gain (50 μL, 60 μL, and 70 μL for days 1, 2, and 3 of feeding, respectively). DSS was dissolved in formula at 0.25, 1, 2, or 3% w/v concentrations.

Neonatal mice fed formula alone, i.e., 0% DSS, served as controls in the characterization of this model to avoid maternal separation as a confounding factor. Unlike breastfed mice, formula-fed mice experience isolation from their mothers, which has been shown to increase stress and anxiety and negatively impacts neurodevelopment [17]. To confirm differences in development between these two groups, control pups were compared to pups nursed by their dams. Breastfed mice demonstrated an increased weight gain as well as differences in cytokine and chemokine profile (Supplemental Figure 1) than formula-fed mice.

Animals were monitored every three hours for signs of NEC (rectal bleeding, lethargy, and abdominal distension). Body weight was recorded every 12 hours, beginning at the initiation of feeds (0 hours). A clinical sickness score (CSS) was assessed, as described by Zani et al. (2008) [18], every 12 hours. As per IACUC protocol, animals were euthanized if they exhibited apnea, rectal bleeding, lethargy, weight loss > 15% of total body weight per day, or no weight gain for two days. Tissue samples were prepared for analysis if animals were euthanized by these indications, thus shortening the duration of the experiment for these pups. The remaining animals were euthanized after 72 hours and tissues were harvested for histological and immunological studies.

### Gastrointestinal Methods

#### GI Evaluation and Preparation

Intestines were placed in cold PBS following dissection from the mouse abdominal cavity. External bowel scores were determined, as described previously by Zani et al (2008) [18]. Bowels were scored upon color, dilation, and friability/consistency for a maximum sickness score of 9 (3 points per category). Intestines were then prepared for fixing, as described in Bialkowska et al. (2016) [19]. Mesentery and connective tissue were removed. The bowel was flushed with modified Bouin’s fixative (50% ethanol / 5% acetic acid in dH_2_O) and cut at the ileocecal junction. Using the Swiss-rolling technique [19], the bowel was fixed overnight in 10% formaldehyde at room temperature and transferred to cold 0.1M PBS the following day. The bowel was paraffin-embedded and sliced into 5 μm sections for hematoxylin and eosin (H&E) or immunohistochemical staining. The Research Histology Core Laboratory, Department of Pathology at Stony Brook University, performed tissue processing, paraffin embedding, slicing, and H&E staining.

#### Intestinal Immunohistochemistry (IHC)

5 μm intestinal paraffin slides were stained as described by Talmasov et al. (2015) [20]. Sections were deparaffinized in xylene, incubated in 2% hydrogen peroxide in methanol for 30 minutes, rehydrated in an ethanol gradient, and then treated with 10 mM Na citrate buffer, pH 6.0, at 120°C for 10 min in a pressure cooker. The slides were then blocked with 5% bovine serum albumin (BSA) in wash solution (0.01% Tween 20, 1X Tris-buffered PBS (TTBS)) and incubated with primary antibodies (Table 1; anti-Ki67, anti-cleaved caspase-3) overnight at 4°C. Slides were washed with TTBS, incubated with a secondary antibody horseradish-peroxidase (HRP) probe from the MACH 3 Rabbit HRP Polymer Detection kit (Biocare Medical, Pacheco, CA), washed with TTBS again, and treated with a tertiary antibody red HRP polymer from the same kit. After the final TTBS washes, color was developed using the Betazoid DAB Chromogen kit (Biocare Medical, Pacheco, CA), and counterstained with hematoxylin solution, Gill (Sigma-Aldrich, St Louis, MO). Slides were dehydrated using alcohol and xylene gradient before mounting.

**Table 1.** Primary (1°) antibodies used for IHC of intestine (paraffin) and brain (frozen) slices.

| 1° antibody | Target | Species | Dilution | Company | Catalog # |
| --- | --- | --- | --- | --- | --- |
| ***Intestine (paraffin)*** |  |  |  |  |  |
| Kiel 67 (Ki-67) | Proliferating cells | Rabbit | 1:200 | Biocare Medical | CRM325B |
| Cleaved caspase-3 (CC3) | Apoptotic cells | Rabbit | 1:400 | Cell Signaling | Ab#9661 |
| ***Brain (frozen)*** |  |  |  |  |  |
| Neuronal nuclear protein (NeuN) | Mature neurons | Mouse | 1:100 | MilliporeSigma | MAB377A5 |
| Ionized calcium binding adaptor molecule-1 (IBA1) | Microglia, macrophages | Rabbit | 1:500 | Wako | 019-19741 |
| Primary antibodies used for immunohistochemical staining of either paraffinized intestinal slices or fixed frozen brain slices (see *Methods*). For the intestines, Ki-67 and CC3 were used independently in different slices, whereas, for the brain, NeuN and IBA1 were co-stained, with DAPI, in the same slices. |  |  |  |  |  |

#### Slide Scanner Light Microscopy

Stained intestinal paraffin images were acquired on an Olympus VS120 Virtual Slide Microscope (Olympus Corporation, Japan) at 20x magnification. Images were processed using Olympus Viewer software, and regions of maximal necrosis or proliferation (based on Ki-67 staining) were saved at 200x magnification (10x zoom of the original scan taken at 20x magnification). These 200x magnification small intestinal images were selected for cell count analysis.

#### Analysis of Intestinal Tissue

H&E-stained intestinal sections were analyzed as per Tanner, et al. (2015) and Liu, et al. (2022) [21, 22]. A minimum of six animals were used for each condition for enterocolitis grading. Intestinal pathology was scored based on observation of the region with the worst lesion. Briefly, the scores were as follows: 0 for intact villi and crypts, 1 for superficial epithelial sloughing, 2 for mild villous or crypt necrosis, 3 for complete villous or crypt necrosis, and 4 for transmural necrosis [21]. For Ki-67 counts, 4 to 10 crypts were identified per 200x magnification image (3 high power fields were analyzed per animal). Ki-67-positive or caspase-positive cells, if present, were counted for each crypt. A minimum of three animals were analyzed for each condition for Ki-67 staining.

### Cytokine and Chemokine Preparation and Analysis

#### Tissue Preparation for Cytokine and Chemokine Measurements

Organ tissue samples (liver and cortex) were collected into sterile microcentrifuge tubes and weighed before being homogenized in sterile endotoxin-free saline using sterile 2.3 mm zirconia/silica beads (Biospec Products Inc.; Bartlesville, OK), as previously described by Speer, et al. (2020) [23]. Samples were then centrifuged at 13,000 g for 10 min at 4°C. Supernatants were harvested and stored at - 80°C for subsequent cytokine and chemokine measurements.

#### Cytokine and Chemokine Analysis

Concentrations of 16 cytokines and chemokines (IL-1α, IL-1β, IL-2, IL-6, IL-10, IL-12(p40), IL-12(p70), IL-17, CXCL1, CCL2, CCL3, CCL4, G-CSF, GM-CSF, IFN-γ, TNF-α) in tissue homogenate supernatants were measured with Bio-Plex Pro magnetic multiplex assays (Bio-Rad; Hercules, CA) and analyzed on the Bio-Plex 200 system with Bio-Plex Manager 5.0 software (Bio-Rad). Results were expressed as pg per mL (plasma) or pg/mg protein (liver, cortex) cytokine or chemokine concentration for supernatants of organ tissue homogenates. Duplicate technical replicates were used for all immunological studies. Protein concentrations in supernatant tissue homogenate samples were determined with the Bradford method (Bio-Rad Laboratories; Richmond, CA) and measured on a Spectramax 190 Plate Reader (Molecular Devices LLC; San Jose, CA).

### Brain Analysis Methods

#### Brain Immunofluorescence

Whole brains were dissected and fixed in 4% paraformaldehyde/phosphate buffer (PFA/PB) at 4°C overnight and then moved to PBS with 30% sucrose/phosphate buffered (PB). After removing the cerebellum, brains were embedded in an optimal cutting temperature (OCT) compound (Fisher Scientific, Hampton, NH). 20 μm sections were made using a cryostat (Leica, Germany). The sections were washed in 0.3% Triton-X100/PBS three times, blocked with 10% goat serum in 0.3% Triton-X100/PBS, and incubated with primary antibodies (Table 1; anti-IBA-1, anti-NeuN conjugated to Alexa-Fluor 555) overnight at 4°C. Next, sections were washed three times in 0.3% Triton-X100/PBS and treated with the secondary antibody goat anti-rabbit Alexa Fluor 488 (ThermoFisher, Waltham, MA; 1:1000) overnight at room temperature. After three washes in 0.3% Triton-X100/PBS, the slides were mounted in DAPI Fluoromount (SouthernBiotech, Birmingham, AL), left to dry overnight, and sealed.

#### Confocal Microscopy and Image Acquisition

Images were captured with an Olympus FV1000 confocal microscope (Olympus Corporation, Japan) with 30x (silicon immersion) and 60x (oil immersion) lenses to image the CA1 hippocampal region of stained sections. 30x magnification images were taken first for neural cell counts. Subsequently, to obtain higher resolution visualization of microglia for Sholl analysis, regions on the 30x magnification images were chosen using a random number generator, to avoid selection bias, for reimaging at 60x magnification. Z-stack images (18-22 stacks, each 1 μm thick) were captured using Olympus FluoView software. Images were subsequently processed and analyzed with Fiji is Just ImageJ (Fiji) software [24]. ***Image Analysis***

All experimenters were blinded to the treatment conditions during image analysis. At least two independent individuals analyzed identical samples. The results were compared to ensure consistency of analysis, and final counts were averaged between two or more individuals. IBA-1 and DAPI co-stained cells were identified as microglia, whereas neurons were identified by their NeuN and DAPI co-staining. DAPI, IBA-1, and NeuN cells were counted for each high-powered field. Cell proportions per image were calculated by dividing the number of microglia or neurons by the number of DAPI-positive cells. Additionally, microglia morphology was analyzed for branching complexity via Sholl analysis [25]. All microglia dendrites (60x magnification) were manually traced and skeletonized. The branching complexity of the skeletonized cells was assessed using the Sholl Analysis plugin for Fiji software (5 μm starting radius, 70 μm ending radius, and 5 μm step size) [26].

### Statistical Analysis

Statistical analysis and graphing of results were performed using Excel and GraphPad Prism software v. 9 and 10 (GraphPad Software; San Diego, CA). P-values less than 0.05 were considered statistically significant. A survival curve log-rank test was performed for mortality. Two-way analysis of variance (ANOVA) tests was used for weight data, behavior scores, intestinal pathology, and Sholl analysis of microglia. One-way ANOVAs were employed for bowel scores and all cell counts (both for the intestines and in the brain). All ANOVA tests were verified by a post-hoc Tukey analysis for all data pair comparisons. Simple linear regression analysis was also conducted for external bowel scores and intestinal cellular proliferation counts. For cytokine and chemokine results, log-transformations, using the equation y = log(y), were first applied to meet the normality assumptions as indicated. Then a simple linear regression model was used to fit the outcome values.

## Results

Enteral administration of formula supplemented with 3% dextran sodium sulfate (DSS), an osmotic agent, to neonatal mice elicited severe enterocolitis characteristic of NEC [15]. Taking advantage of this general approach, we fed neonatal mice either 0% (control) or increasing concentrations of DSS (0.25%, 1%, 2%, or 3%) for 72 hours to determine if we could titrate the severity of NEC induced to mimic NEC stages seen in neonatal human patients. We first assessed the clinical characteristics, such as mortality, weight changes, and sickness scores.

### Increasing DSS supplementation increases mortality rate, diminishes weight gain, and results in earlier abnormal clinical sickness scores

Approximately 50% of all experimental mice survived the feeding protocol. Mice either died between feeds or required euthanasia, as per the IACUC protocol (*see Methods*). Mice in the 0%, 0.25%, and 1% DSS groups had comparable Kaplan-Meier survival curves (Figure 1A), which all differed significantly from survival curves in the 2% and 3% DSS groups (*p <* 0.01, pairwise Kaplan-Meier survival analysis between the three low DSS groups and the two high DSS groups, survival curve log-rank). All mice in the 2% DSS group died or were euthanized before 72 hours, and mice in the 3% DSS group died or were euthanized before 48 hours (*p <* 0.01, survival curve log-rank, for 2% versus 3% DSS).

**Figure 1.**
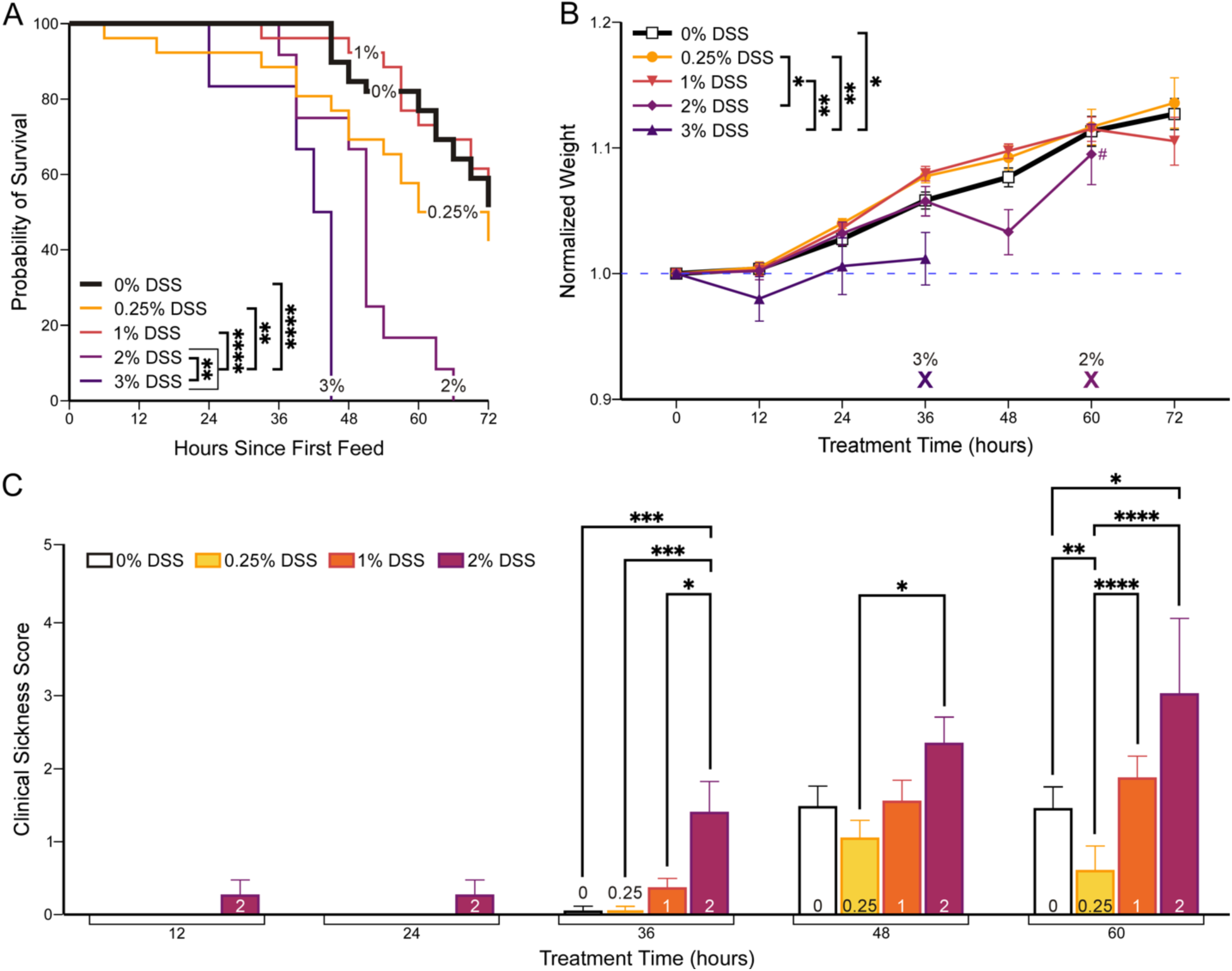
Survival, weight gain, and clinical sickness scores (CSS) correlate with dextran sodium sulfate (DSS) concentration. Starting on postnatal day 3, mice were fed either 0% (control) or increasing concentrations of DSS (0.25%, 1%, 2%, or 3%) for 72 hours (see Methods). **(A)** Kaplan-Meier survival curve. Mice fed high concentrations of DSS (2% & 3%) had a shorter lifespan compared to lower concentrations. Pairwise survival analysis between high DSS and low DSS groups, *p* < 0.0001 (Supplemental Table 1). Number of mice: 0%, 40; 0.25%, 26; 1%, 26; 2%, 12; 3%, 6. **(B)** Normalized weight trends during protocol. **X** indicates the time point at which all mice died or were euthanized within that experimental group. **#** at 60 hours where only 2 mice have survived in 2% DSS (n: 12 → 2). Two-way ANOVA with Tukey’s post-hoc, *p* < 0.0001 (Supplemental Tables 2 and 3). Number of mice same as in 1A. **(C)** CSS increases (worsens) more rapidly in animals exposed to higher DSS concentrations (see Supplemental Figure 2 for scores per category). Two-way ANOVA with Tukey’s post-hoc, *p* < 0.0001 (Supplemental Tables 4 and 5). Number of mice: 0%, 29; 0.25%, 26; 1% 23; 2%, 7. 3% DSS data not shown because mice were too sick to be assessed. Data presented as mean ± SEM. \**p* < 0.05, \*\**p* < 0.01, \*\*\**p* < 0.001, \*\*\*\**p* < 0.0001.

To determine if DSS concentration would affect growth during the feeding period, we weighed mice every 12 hours. Higher concentrations of DSS supplementation (2% and 3%) resulted in overall weight loss, whereas mice fed with lower DSS concentrations (0%, 0.25%, 1%) increased their weight (Figure 1B). The final mean normalized weights at 72 hours are summarized in Supplemental Table 2. There was no statistically significant difference in overall weight gain among low DSS concentration groups (0%, 0.25%, 1%) (*p =* 0.354, two-way ANOVA with Tukey’s post-hoc). Between the 48- and 60-hour time points, only 20% of the 2% DSS mice survived, which explains the sudden deviation in the mean normalized weight. The 3% DSS mice deteriorated rapidly; therefore, we omitted 3% DSS-fed mice from subsequent studies and used 2% DSS to represent severe enterocolitis.

Throughout the feeding protocol, we used a clinical sickness score (CSS) [18] to assess neonatal mouse behavior and health. Mice in the 2% DSS group began exhibiting signs of illness – concerning appearance, activity, and response to touch – as early as 12 hours of DSS administration; the CSS in these mice continued to increase over time (Figure 1C). In contrast, mice exposed to lower concentrations of DSS (0%, 0.25%, 1%) began exhibiting abnormal behavior only after 36 hours of feeding. 2% DSS-fed mice demonstrated significantly higher clinical sickness scores compared to all other DSS conditions after 36 hours (*p* < 0.05, two-way ANOVA with Tukey’s post-hoc, Supplemental Table 5). Overall, increased DSS supplementation resulted in significantly decreased survival, decreased weight gain, and worse clinical sickness scores in mice.

### DSS feeds induced enterocolitis in neonatal mice

NEC causes intestinal inflammation, inflammatory cell recruitment, and villous sloughing. This process progresses through the muscularis layer and can eventually result in transmural necrosis and perforation [21]. After three days of feeding, we evaluated histology (Figure 2A) and gross pathology of the intestines to determine if increasing concentrations of DSS resulted in worsening NEC. The external bowel score assesses gut color, dilation, and consistency or friability [18]. We found that the external bowel score was significantly correlated with increasing DSS concentrations (slope: 2.41 with *p* < 0.0001, Figure 2B, Supplemental Figure 3, Supplemental Table 6 & 7).

**Figure 2.**
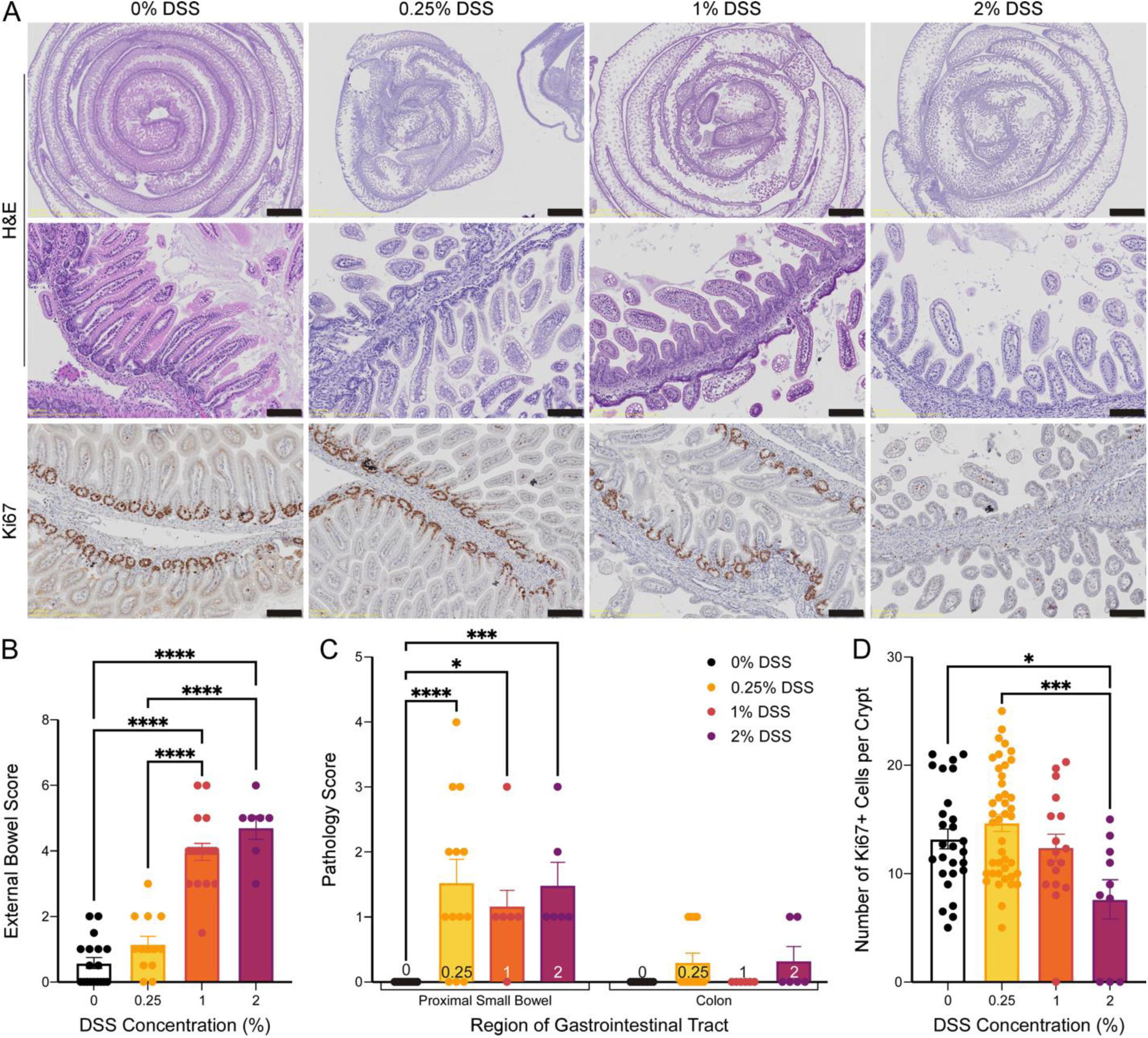
Enterocolitis severity increases with DSS concentration. Increased DSS concentration led to increased external bowel score, disruption of intestinal morphology, and decreased Ki-67+ cells. **(A)** Representative images of hematoxylin- and eosin-stained (H&E) intestinal samples imaged at 10x (1^st^ row, scale 1 mm) or at 200x magnification (2^nd^ row, scale 100 µm) or of Ki-67 immunohistochemistry at 200x magnification (3^rd^ row, scale 100 µm). **(B)** External bowel severity scores are higher (worse) at increasing DSS concentrations, 1 & 2% (see Supplemental Figure 3 for scores per category). One-way ANOVA with Tukey’s post-hoc, *p* < 0.0001 (Supplemental Tables 6 and 7). Number of mice: 0%, 18; 0.25%, 13; 1%, 18; 2%, 7. **(C)** Enterocolitis pathology, which was only present in DSS groups, was primarily localized to the small intestine. Two-way ANOVA with Tukey’s post-hoc, *p* < 0.0001 (Supplemental Table 8). Number of mice: 0%, 7; 0.25%, 7; 1%, 6; 2%, 3. **(D)** Cellular proliferation, as measured by number of Ki67-positive cells within small intestinal crypts, decreased significantly in 2% DSS. One-way ANOVA with Tukey’s post-hoc, *p* < 0.0001 (Supplemental Tables 9 and 10). Number of crypts: 0%, 27; 0.25%, 42; 1%, 16; 2%, 7. Data presented as mean ± SEM. \**p* < 0.05, \*\**p* < 0.01, \*\*\*\**p* < 0.0001.

Complementing the gross pathology, H&E staining of DSS-treated intestines (Figure 2A) also indicated the presence of enterocolitis with derangement of villous organization, intestinal dilatation with patchy necrosis and perforation. Consistent with previous literature [15], enteral DSS administration primarily caused enterocolitis in this newborn mouse model (Figure 2B and C), which contrasts with the colitis observed in older mice treated with DSS [27]. We scored the intestinal pathology based on the worst pathology observed in the slide. All mice exposed to DSS demonstrated enterocolitis. Intestines from mice fed 0.25%, 1% and 2% DSS scored significantly worse than those from 0% DSS-fed mice (*p <* 0.001, *p* < 0.05 and *p =* 0.007, respectively). Animals administered 0.25% and 2% DSS displayed mild colitis, which was not significantly different from control. Following these findings, we performed immunohistochemistry to determine if enterocolitis was associated with cellular apoptosis or proliferation abnormalities.

### Increased DSS exposure suppresses intestinal cell proliferation without promoting apoptosis

We assessed the intestines of DSS-exposed and formula-fed (control) mice for the presence of cellular apoptosis or proliferation by staining for cleaved caspase-3 (CC3) (data not shown) or Ki-67 (Figure 2A, bottom row), respectively. For all experimental groups, small intestinal crypts did not show any positive staining for CC3 (data not shown). In contrast, Ki-67 positive (Ki-67+) staining of small intestinal crypts in 2% DSS-fed mice was significantly decreased compared to the staining in 0% or 0.25% DSS-fed pups (Figure 2D). Simple linear regression analysis of the Ki-67+ cell counts demonstrates that cell proliferation was negatively correlated with DSS concentration (slope: -2.94, *p* = 0.0009).

### Higher DSS concentrations in feeds led to hepatic and systemic inflammation

Overall, we observed that higher concentrations of DSS resulted in worsening clinical signs, intestinal pathology, and diminished intestinal cell proliferation. NEC causes local intestinal inflammation accompanied by cytokine and chemokine production that propagates to a systemic response [28]. We therefore measured cytokine and chemokine levels in the liver to determine if the levels of these inflammatory mediators correlate with the DSS concentration and the severity of enterocolitis.

Specifically, we measured 16 cytokines and chemokines (IL-1α, IL-1β, IL-2, IL-6, IL-10, IL-12(p40), IL-12(p70), IL-17, IFN-γ, TNF-α, G-CSF, GM-CSF, CCL2, CCL3, CCL4, CXCL1) that are involved in sepsis from liver samples. Since the liver is the first major filtration organ downstream of the intestines, we used liver data as a measure of gastrointestinal cytokine and chemokine production. Notably, all cytokine concentrations in liver tissue positively correlated with increasing supplementation of DSS (Figure 3). Chemokine production induces the release and recruitment of immune cells to the peripheral tissues; thus, chemokine levels also correlated with DSS supplementation (Figure 4). We performed the same analysis on blood plasma; several cytokine and chemokine levels were below the threshold of detection. In plasma, concentrations of IL-12, G-CSF, CXCL1, CCL4 and IL-10 correlated with increasing DSS supplementation (Figure 5).

**Figure 3.**
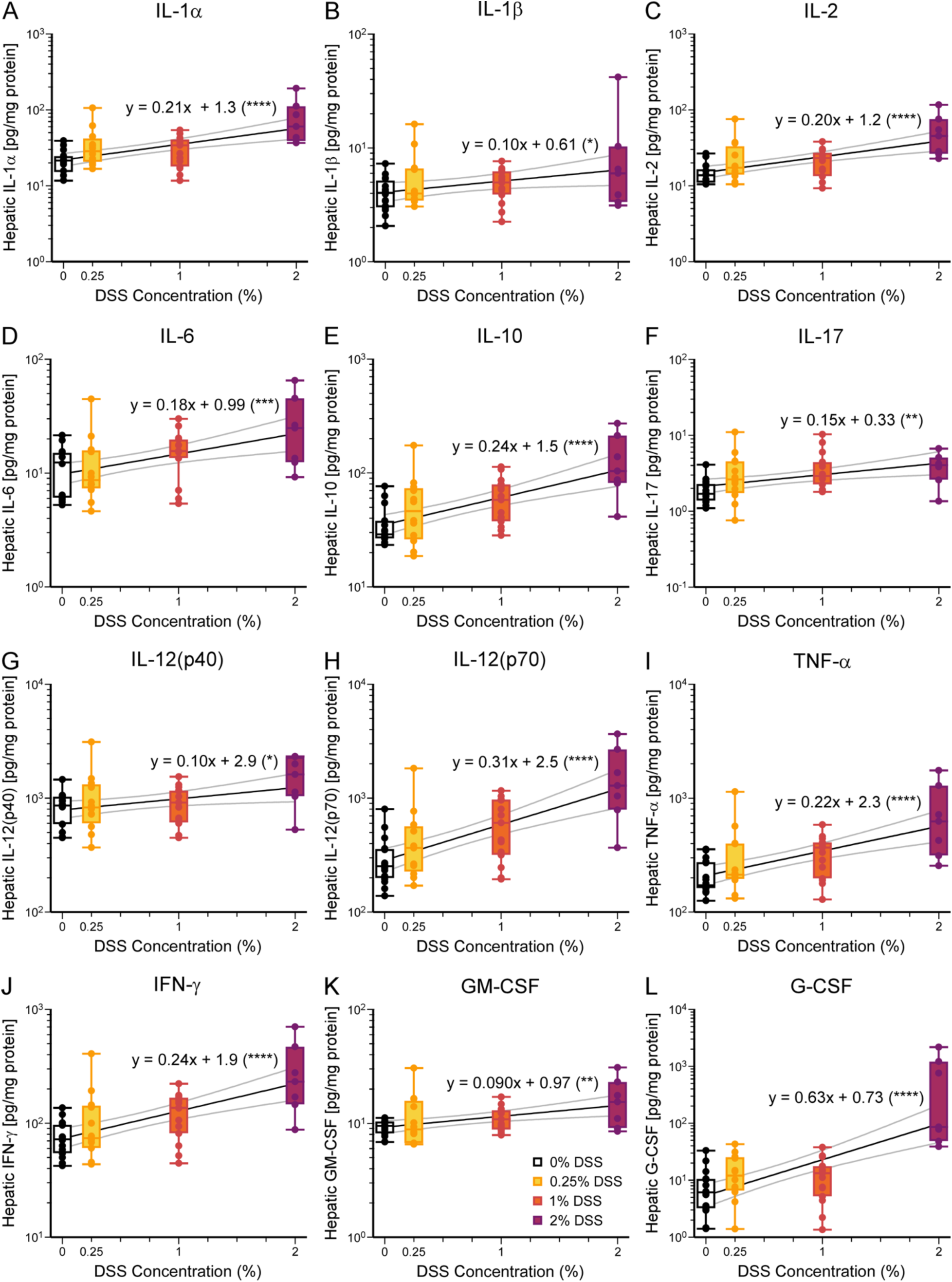
Concentrations of all tested hepatic cytokines positively correlate with DSS concentration. **(A-H)** In the liver, cytokine concentrations, indicative of systemic inflammation, are all positively correlated with DSS concentrations. Cytokines tested include: **(A)** IL-1α, *p* < 0.0001, **(B)** IL-1β, *p* = 0.033, **(C)** IL-2, *p* < 0.0001, **(D)** IL-6, *p* = 0.0009, **(E)** IL-10, *p* < 0.0001, **(F)** IL-17, *p* = 0.0035, **(G)** IL-12(p40), *p* = 0.020, **(H)** IL-12(p70), *p* < 0.0001, **(I)** TNF-α, *p* < 0.0001, **(J)** IFN-γ, *p* < 0.0001, **(K)** GM-CSF, *p* = 0.0024, and **(L)** G-CSF, *p* <0.0001. Simple linear regression with log-transformation of y values performed (see Supplemental Table 11). Data presented as boxplots showing min-max. Slope and y-intercept with confidence intervals are plotted. \**p* < 0.05, \*\*\**p* < 0.001, \*\*\*\**p* < 0.0001. Number of mice: 0%, 16; 0.25%, 12; 1%, 15; 2%, 7. *p*-values calculated for t statistic of slope (coefficient of DSS concentration).

**Figure 4.**
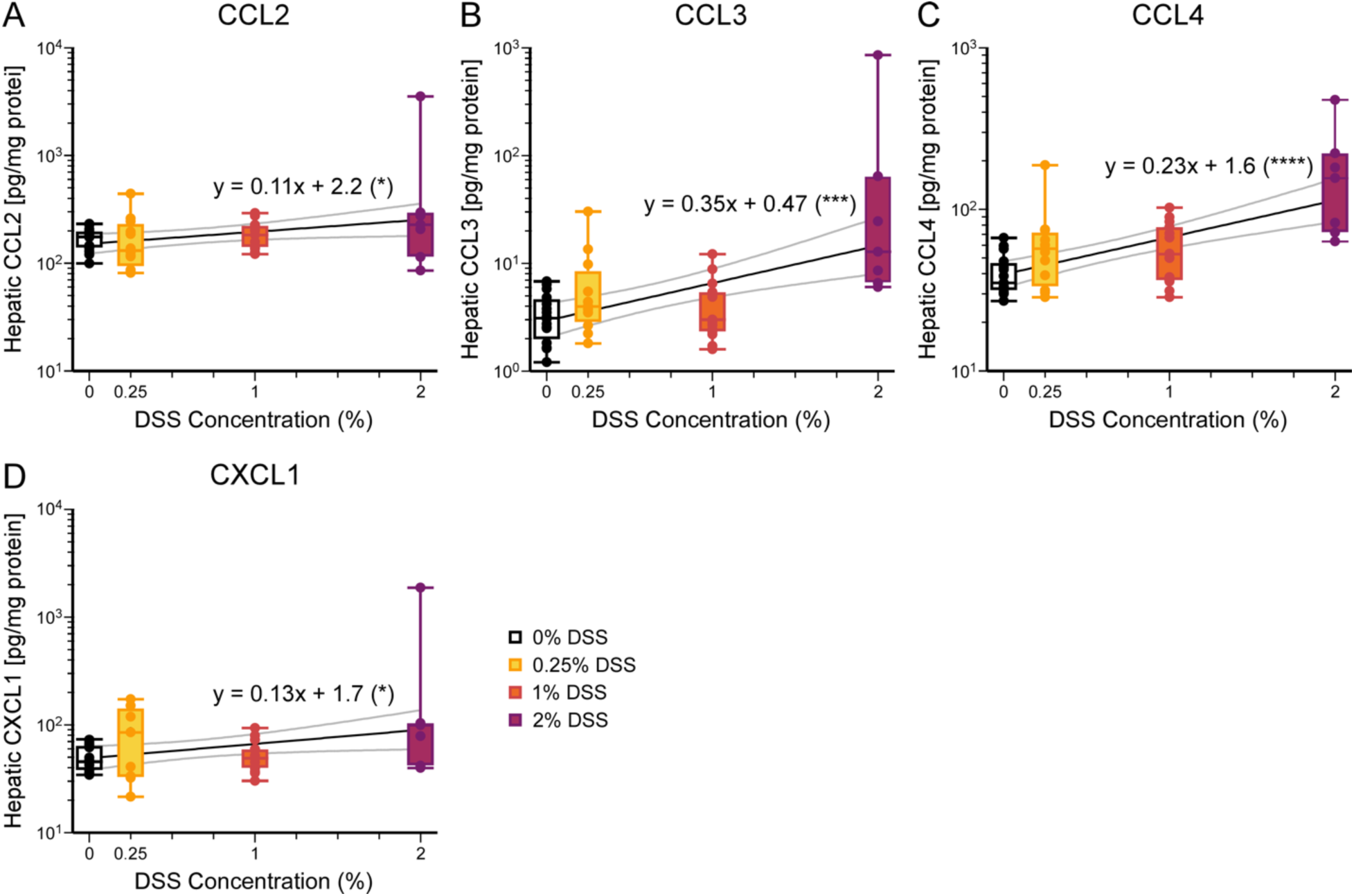
Concentrations of hepatic chemokines positively correlate with DSS concentration. **(A-H)** In the liver, chemokine concentrations are positively correlated with DSS concentrations. Chemokines tested include: **(A)** CCL2, *p* = 0.027, **(B)** CCL3, *p* = 0.0002, **(C)** CCL4, *p* < 0.0001, and **(D)** CXCL1, *p* = 0.031 (see Supplemental Table 11). Results displayed and analyzed as in Figure 3. \**p* < 0.05, \*\*\**p* < 0.001, \*\*\*\**p* < 0.0001. Number of mice: 0%, 16; 0.25%, 12; 1%, 15; 2%, 7.

**Figure 5.**
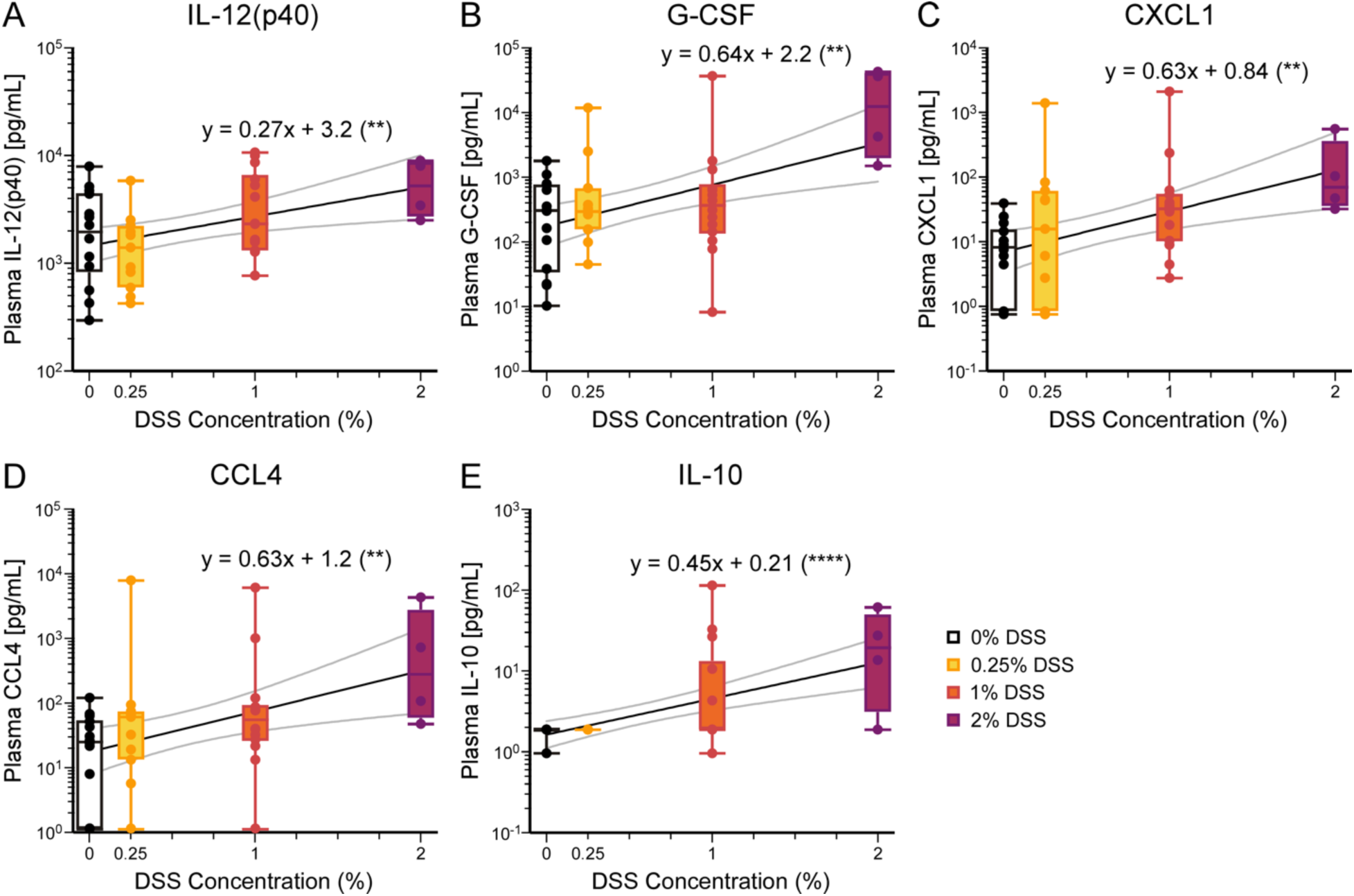
Some blood plasma cytokine and chemokine concentrations correlate with DSS concentrations. **(A-E)** In blood plasma, concentrations of **(A)** IL-12(p40), *p* = 0.0066, **(B)** G-CSF, *p* = 0.0016, **(C)** CXCL1, *p* = 0.0016, **(D)** CCL4, *p* = 0.0045, and **(E)** IL-10, *p* < 0.0001, are positively correlated with DSS concentrations (see Supplemental Table 12 and Supplemental Figure 4). Results displayed and analyzed as in Figure 3. \**p* < 0.05, \*\**p* < 0.01. Number of mice: 0%, 14; 0.25%, 11; 1%, 14; 2%, 4.

In summary, increasing concentrations of DSS resulted in worsening clinical phenotypes, worsening enterocolitis, and increasing inflammation in liver tissue and plasma, as assayed by cytokines and chemokines. Given this strong association between the DSS concentration and NEC severity, we next used this model to study the neurological sequelae of NEC in mice.

### NEC induces neuroinflammation

Neurodevelopmental impairment is a significant long-term morbidity associated with NEC [6]. Even patients who have experienced milder forms of NEC are at a higher risk of cognitive disability compared to age-matched premature infants without NEC [29]. We, therefore, measured cytokine and chemokine concentrations in the brain of DSS-treated mice.

We measured the same array of cytokines and chemokines as for hepatic and systemic inflammation in brain tissue (Figure 6 & Supplemental Figure 5). Overall, the concentrations of cytokines and chemokines measured in the brain were lower than those measured in the liver. Two cytokines (IL-2, G-CSF) and one chemokine (CXCL-1) increased with DSS concentration (Figure 6). Therefore, increasing the severity of NEC induced by DSS resulted in increasing concentrations of a wide array of inflammatory cytokines and chemokines in the liver and plasma but only a small subset in the brain.

**Figure 6.**
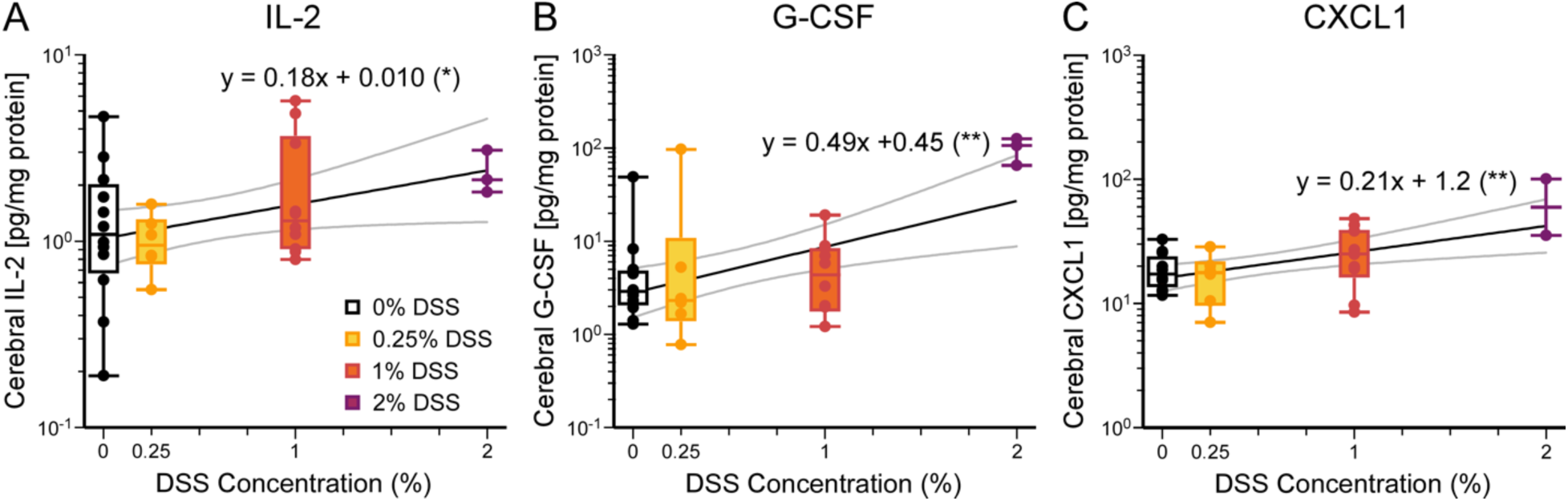
Cerebral cytokines IL-2 and G-CSF and chemokine CXCL1 concentrations positively correlate with DSS concentrations. **(A-C)** In the brain, concentrations of **(A)** IL-2, *p* = 0.043, **(B)** G-CSF, *p* = 0.0030, and **(C)** CXCL1, *p* = 0.0035, are positively correlated with DSS concentrations (see Supplemental Table 13 and Supplemental Figures 5 and 6). Results displayed and analyzed as in Figure 3. \**p* < 0.05, \*\**p* < 0.01. Number of mice: 0%, 12; 0.25%, 6; 1%, 10; 2%, 3.

### NEC does not affect the number of neurons or microglia but increases microglia activation in the CA1 region of the hippocampus

We next determined the impact of graded NEC on central nervous system anatomy, specifically for neurons and microglia, the resident immune cells of the central nervous system. (Figure 7). There was a slight increase in neuron proportion between 1% to 2% DSS supplemented mice (*p* = 0.027, One-way ANOVA with Tukey’s post-hoc); otherwise, cell counts across groups were comparable. For microglia, there was a statistically significant difference only between 0.25% and 1% DSS conditions (0.25 vs. 1%, *p =* 0.018; 0.25 vs. 2%, p = 0.068, One-way ANOVA with Tukey’s post-hoc).

**Figure 7.**
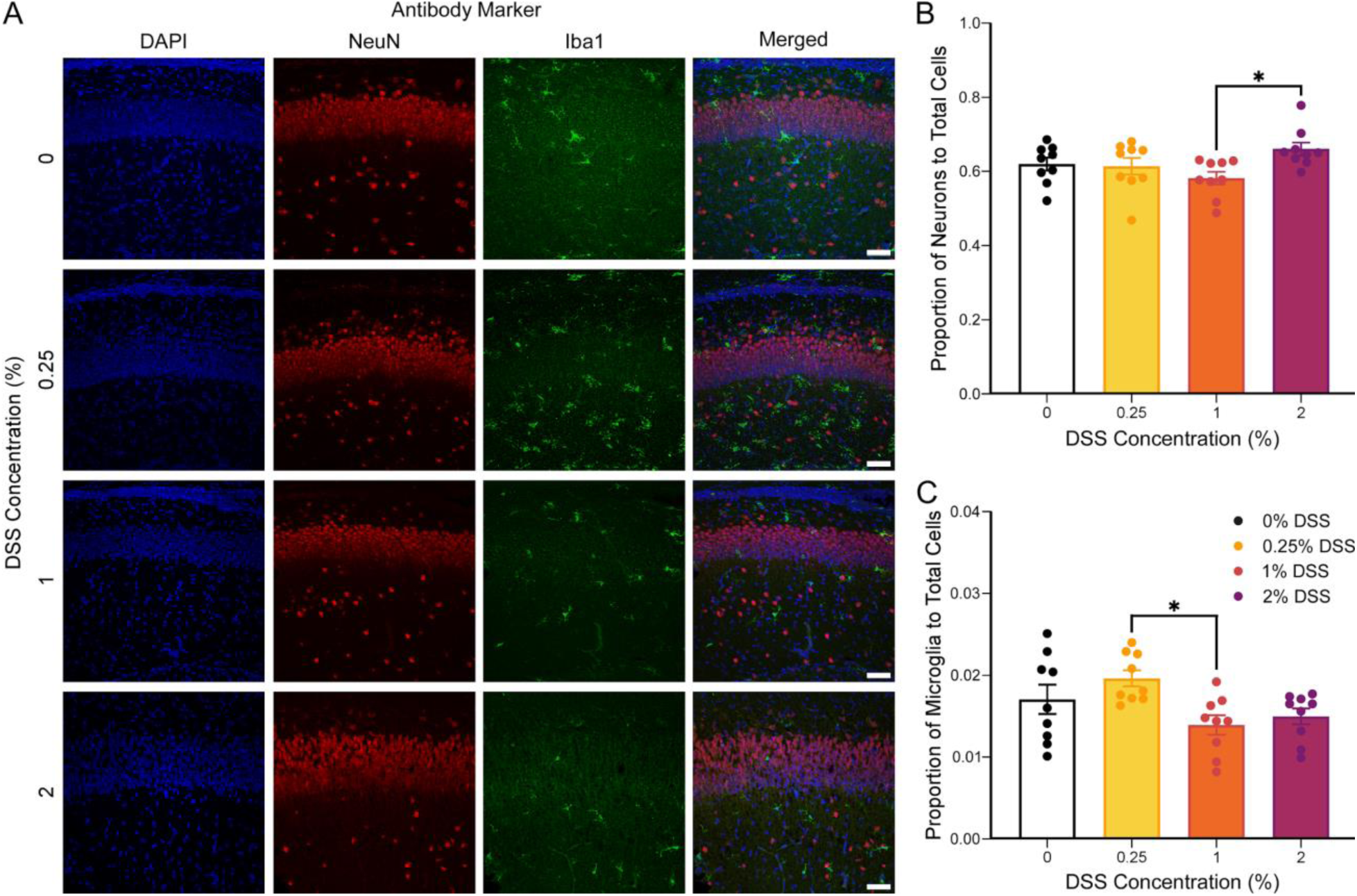
DSS-induced NEC does not change neuron or microglia proportions in the CA1 hippocampal region. **(A)** Representative immunohistochemical images (30x magnification) of the CA1 hippocampal region. Slices are stained blue for DAPI, red for neurons, and green for microglia. Scale bar = 50μm. **(B-C)** The proportion of **(B)** neurons, *p* = 0.046, and **(C)** microglia, *p* = 0.018, out of all cells in the CA1 region is comparable across all DSS concentrations (see Supplemental Figure 7 for raw counts). A significant difference was only found when comparing the proportions of neurons between mice fed 1% and 2% DSS, *p* = 0.027 (Supplemental Table 14), and when comparing the proportions of microglia between mice fed 0.25% and 1% DSS, *p* = 0.018 (Supplemental Table 15). One-way ANOVA with Tukey’s post-hoc. Data presented as mean ± SEM. \**p* < 0.05. n = 9 immunohistochemical images for all experimental groups.

Microglia retract their filopodia upon activation and assume a more amoeboid shape [30, 31]. To assess microglia morphology, we applied Sholl analysis to 60x images of IBA-1 positive cells (Figure 8A and 8B) [25]. Branching intersections peaked at 10 – 15 microns from the cell soma and tapered at 70 microns (Figure 8C, Table 2). Microglia from control (0% DSS) mice demonstrated significantly greater branching patterns compared to all mice with DSS supplementation (0.25, 1, 2%) (*p <* 0.0001, Two-way ANOVA with Tukey’s post-hoc). Given that the branching complexity of microglia is inversely related to their activation level [31], these results indicate that DSS-fed mice displayed more activated microglia than those in control (0% DSS-fed) mice.

**Figure 8.**
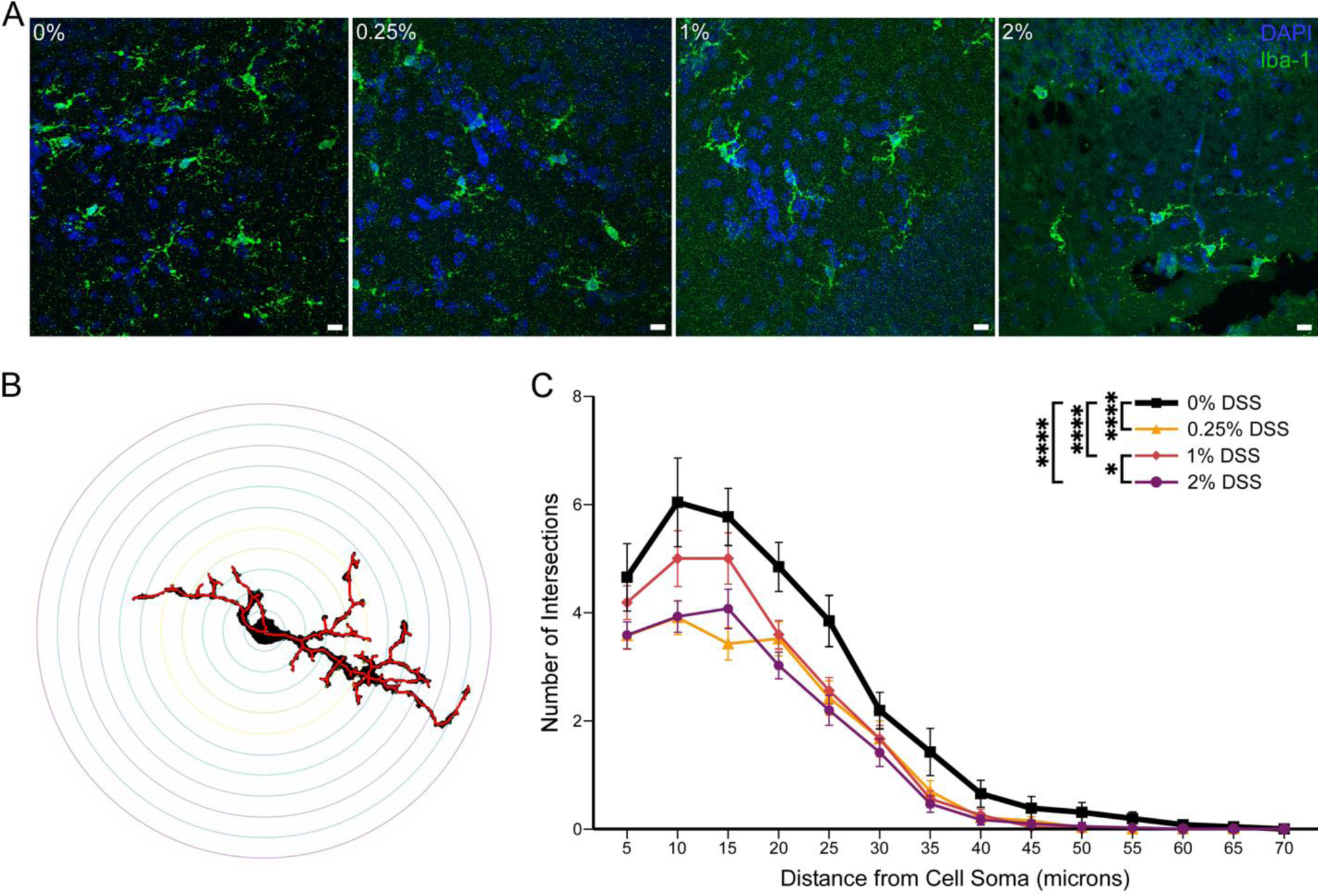
DSS supplementation at any concentration activates microglia in the CA1 hippocampal region. **(A)** Representative immunohistochemical images (60x magnification) of microglia in the CA1 hippocampal region. Slices are stained blue for DAPI and green for microglia. Scale bar = 10 μm. **(B)** Representative Sholl analysis tracing, with skeletonized trace in red, of a microglial cell identified by Iba-1 (0% DSS). **(C)** Sholl analysis. All DSS concentrations showed a reduced number of branching intersections (more activation) than the 0% DSS-fed mice. Two-way ANOVA with Tukey’s post-hoc, *p* < 0.0001 (see Table 2). Number of microglia: 0%, 26; 0.25%, 33; 1%, 27; 2%, 41. Data presented as mean ± SEM. \**p* < 0.05, \*\*\*\**p* < 0.0001.

**Table 2.** Sholl analysis comparisons at different distances from microglial cell soma.

| Comparison | Distance from Cell Soma (microns) |  |  |  |  |  |  |  | Overall |
| --- | --- | --- | --- | --- | --- | --- | --- | --- | --- |
|  | 5 | 10 | 15 | 20 | 25 | 30 | 35 | 40 |  |
| 0% vs 0.25% DSS | **0.021** | **<0.0001** | **<0.0001** | **0.0022** | **0.0009** | 0.50 | 0.21 | 0.64 | **<0.0001** |
| 0% vs 1% DSS | 0.63 | **0.041** | 0.20 | **0.0078** | **0.0056** | 0.54 | 0.12 | 0.75 | **<0.0001** |
| 0% vs 2% DSS | **0.015** | **<0.0001** | **<0.0001** | **<0.0001** | **<0.0001** | 0.13 | **0.037** | 0.53 | **<0.0001** |
| 0.25% vs 1% DSS | 0.35 | **0.017** | **0.0001** | >0.99 | 0.98 | >0.99 | 0.98 | >0.99 | 0.089 |
| 0.25% vs 2% DSS | >0.99 | >0.99 | 0.21 | 0.46 | 0.90 | 0.87 | 0.90 | >0.99 | 0.97 |
| 1% vs 2% DSS | 0.33 | **0.013** | **0.044** | 0.38 | 0.74 | 0.89 | 0.99 | 0.99 | **0.020** |

## Discussion

Necrotizing enterocolitis is a major cause of acute and long-term morbidity in premature infants [2]. Novel approaches are needed to understand this multifactorial disease process [32, 33]. In patients, NEC causes symptoms of increasing severity; patients develop abdominal distension and mild enterocolitis, which can progress to fulminant sepsis, intestinal necrosis, and death. We have characterized a neonatal mouse model where NEC severity can be prospectively controlled, as demonstrated by clinical and behavioral data, intestinal pathology, and cytokine and chemokine measurements. We show that enterocolitis induces neuroinflammation by causing microglial activation and increased expression of cerebral cytokines and chemokines. These data corroborate clinical studies where NEC patients demonstrate an increased risk of neurocognitive impairment [6].

### A Graded Mouse Model of NEC

DSS administration to adult mice primarily leads to colitis [27], whereas in neonatal mice and rats, it causes small and large bowel inflammation [15, 34]. We supplemented enteral feeds with different concentrations of DSS – 0.25%, 1%, 2% and 3% DSS – to create a graded model of NEC. To avoid maternal separation as a confounding factor [17], mice fed formula (0% DSS) served as control for our experiments.

Although Ginzel et al. used 3% DSS supplementation to induce NEC in neonatal mice [15], we found that 3% DSS caused significant mortality within 48 hours (Figure 1A). The discrepancy in animal mortality and colitis severity may be due to differences in the mouse sub-strain or even their microbiomes, which have been shown to vary in the same mouse strains from different laboratories [35]. Therefore, we used the 2% DSS concentration group to represent severe enterocolitis because 3% DSS supplemented mice died too soon for adequate experimental analysis.

With increasing DSS supplementation, mortality increased (Figure 1A). Similarly, patients with more severe NEC (i.e., Bell’s class III or surgical NEC), have a higher mortality rate [4]. Fullerton et al. show that 74% of medically managed NEC patients survived to follow-up at 1-2 years of age, whereas only 62% of surgical NEC patients did [29]. Mortality from NEC animal protocols ranges from 0 to 30% [33]. In our experiments, formula-fed mice also had a significant mortality rate, which highlights the difficulty of gavage feeding of neonatal mice (2 grams) who are also stressed by separation from their dams [17].

The weight and clinical sickness score mirror the mortality data, where enterocolitis severity correlated with DSS supplementation (Figure 1B and 1C). The most severe NEC (3% DSS) mice never gained weight, while mild (0.25%) and control (0%) mice steadily gained weight. The 2% and 1% DSS treated mice started to lose weight at the end of the protocol, suggesting that the mice developed delayed enterocolitis. Mice receiving high DSS supplementation (2%) developed worse clinical sickness scores earlier than those exposed to lower concentrations. Hence, we could prospectively control and create a graded NEC model by modulating enteral DSS supplementation.

Consistent with other groups [15, 34], DSS exposure in neonatal mice results in enterocolitis and, to a lesser extent, colitis (Figure 2A-C), which contrasts with DSS’s effects in adult mice [27]. Increased DSS supplementation resulted in worse macroscopic gut assessment scores (bowel distension, bloody stool, and friability), which correlates with NEC severity in other models [18]. Although there were no differences in apoptosis as indicated by cleaved caspase-3 staining, increased DSS supplementation decreased intestinal cell proliferation in a concentration-dependent manner, as shown by Ki-67 staining (Figure 2D). Our institution’s IACUC protocol mandates euthanasia in moribund animals or animals with gross bloody stool; this may have accounted for the absence of caspase staining in our samples and decreased colitis observed, as animals may have been euthanized before developing fulminant colitis. Histological hallmarks of NEC such as villous sloughing and atrophy, necrosis and inflammatory infiltrates were seen on hematoxylin and eosin staining, all consistent with other models of NEC [21, 33, 36].

A critical component of NEC pathogenesis is the initiation and propagation of the systemic inflammatory response [28, 37]. NEC patients have increased levels of cytokines and chemokines in the plasma (IL-1β, IL-6, IL-10, IL-17, TNF-α, CXCL1, CCL2, CCL3, and CCL4) [38–41]. Our study showed that all tested liver cytokines and chemokines were increased with higher DSS concentration, thus worsening NEC. Upon closer examination, the specific pattern of increase varied across the cytokines and chemokines. For example, severe enterocolitis (2% DSS) caused a significant increase in liver G-CSF, suggesting that additional pathways may be recruited, or an additional factor caused acceleration of G-CSF production. In the plasma samples, several cytokines evaluated were below the level of detection, possibly due to cytokine and chemokine concentration fluctuations during the inflammatory process. Other animal models of NEC also demonstrate elevations in IL-6, TNF-α, IFN-γ, IL-1β, CXCL1 systemically and in the brain [12–15, 34, 40]; our study looked at the most extensive panel of cytokines and chemokines to date.

### Implications for the Gut-Brain Axis

Neurodevelopmental delay and sensorimotor disabilities are a major long-term morbidity found in NEC patients [42, 43]. Severe NEC, especially if surgery is required, significantly increases the risk of cognitive and sensorimotor disabilities, and affects patients’ school performance [8]. Nevertheless, premature children who had mild NEC are also at an increased risk for adverse neurodevelopmental outcome when compared to gestational age-matched controls [6].

Systemic cytokines and chemokines induced by NEC are believed to be one of the pathways whereby gut inflammation induces changes within the central nervous system during neurodevelopment [1]. NEC induced inflammatory factors cause breakdown of the blood brain barrier and allow neuroinflammatory pathways and effector cells to be activated. We find that brain levels of IL-2, G-CSF, and CXCL1 are increased with higher DSS concentrations (Figure 4). There are many explanations for the increase of a smaller number of cytokines and chemokines in the brain during NEC. Pro- and anti-inflammatory cytokines have early and late phase components; early phase cytokines such as TNF-α may already be declining when measured in the brain in our NEC model. Alternately, the blood brain barrier may filter circulating cytokines and chemokines and limit the number of cytokines activating neuroinflammation [28]. The brain is a heterogenous organ, so specific regions may have cytokine and chemokine levels below the limit of detection by our methods. Both IL-2 and G-CSF knockout mice have neurodevelopmental deficits, therefore it is intriguing to consider the neurodevelopmental effects elevated levels of IL-2 and G-CSF levels would have in this immature brain [44, 45].

Microglia are a major mediator of neuroinflammation during NEC. Nino et al. have shown that reducing microglia activation by interfering with neuroinflammation pathways decreases white matter lesions and learning deficits in mice who have experienced NEC [11]. Although we did not observe an increase in microglia that has been reported [12, 46], we observed a significant change in microglia activation in all mice exposed to DSS (Figures 5 and 6), which is similar to the result of Sun et al. [14]. NEC causes neuroinflammation by increasing cytokine and chemokine levels and activating microglia.

We did not find a difference in NeuN-positive mature neurons in mice with enterocolitis versus controls (Figure 5). Our microglia and neuron results contrast with other groups, which found that NEC induction decreased mature neurons and increased microglia numbers in the hippocampus, basal ganglia, and cerebral cortex of pigs and mice [13, 34, 47]. However, these studies used different ages, induction protocols, and employed different animal species, where cell development and number may innately differ. The mouse age chosen in this model (postnatal day 3 – 6) mirrors central nervous system development of third trimester human infant, thus allowing us to study NEC’s effect upon premature infants, whereas many models look at a mouse age whose brains are undergoing development of term infants [16].

In our graded NEC model, we can induce and control enterocolitis severity by varying DSS supplementation. Specific cytokine and chemokine levels in liver, plasma, and brain correlate with the amount of DSS supplementation (Figure 4) and, thus, the severity of enterocolitis induced. Microglia activation, as demonstrated through our findings, is an all-or-none phenomenon where even mild enterocolitis (0.25% DSS) induces microglial activation to the same extent as worse NEC (Figure 6). These results raise intriguing questions on how NEC induced changes in brain cytokines, chemokines, and microglia that may disrupt normal brain development (Figure 9). Cytokines have been shown to affect synaptic scaling and circuit whereas IL-2 knockout mice exhibit deficits in hippocampal dependent learning and memory [45, 48, 49]. Microglia participate in refinement of neuronal synaptic circuitry as well as embryonic neurodevelopment. Understanding how cytokine, chemokine and microglia interact and change brain development may help identify targets to prevent neurodevelopmental defects induced by NEC.

**Figure 9:**
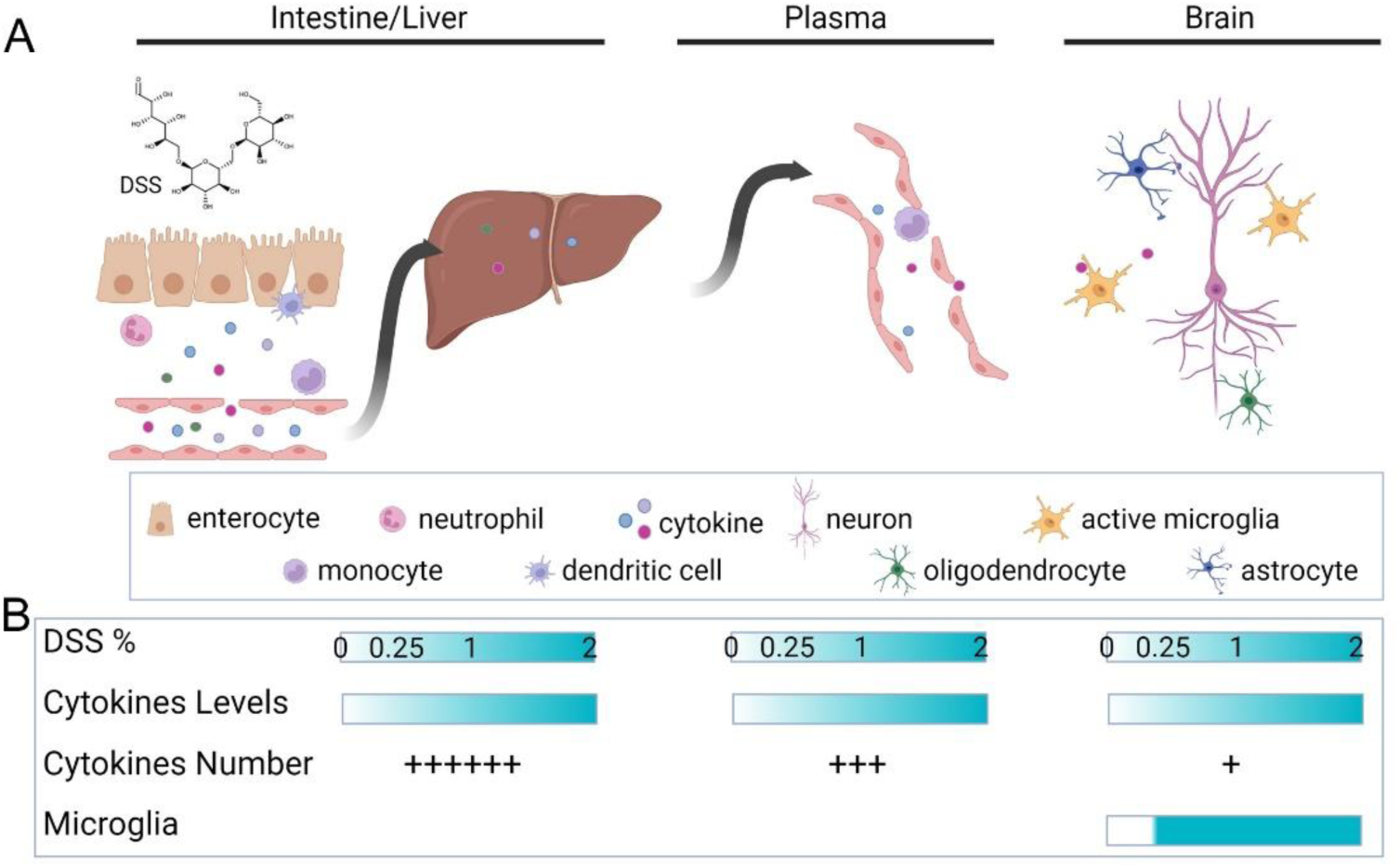
Necrotizing Enterocolitis, cytokines, and neuroinflammation. **(A)** DSS exposure induces intestinal damage and activation of innate immune responses in the neonatal gut with subsequent release of inflammatory cytokines and chemokines into the surrounding tissue. These inflammatory mediators reach the liver via the portal venous system. Systemic circulating inflammatory factors can reach the brain when they cross a leaky blood brain barrier. In the brain, inflammatory cytokines and chemokines can interact with resident immune cells, microglia, or invading peripheral immune cells. These immune cells, cytokines, and chemokines can trigger neuroinflammation directly or indirectly on neurons, oligodendrocytes, and astrocytes. For example, cytokines may activate microglial engulfment of synaptic spines and affect circuit maturation during development. Activation of oligodendrocytes and astrocytes may affect myelination. **(B)** Inflammatory cytokine and chemokine expression increase with rising DSS concentrations and worsening enterocolitis. There number of cytokines whose concentration correlates with DSS concentration decreases as we go from the liver to plasma and, eventually, to the brain. Compared to these patterns in cytokine expression, microglia activation occurs even at low levels of DSS supplementation and mild enterocolitis. Figure created using BioRender.

### Limitations and Considerations

Animal models are crucial for advancing our understanding of the gut-brain axis, and for identifying mechanisms underlying the long-term neurodevelopmental deficits seen in NEC patients [13, 18, 21, 33]. A major consideration when using animal models is the small sample size. By testing several severities of NEC in neonatal mice as well as examining multiple tissues with different experimental modalities, the number of samples was limited. Additionally, these results represent a single time point; future studies would include time course experiments whereby one can study the evolution of mild NEC to severe NEC. Finally, neurodevelopment in adult and adolescent NEC animals can be evaluated with learning and memory paradigms and neuroanatomy.

The unique characteristics and strengths of our DSS rodent NEC model are 1) using a single hyperosmolar agent, DSS, and 2) the mouse age. Despite differences in NEC induction technique, we see similar histopathological changes and cytokine and chemokine changes than those observed in other animal models and in human NEC patients. Furthermore, the earlier age of NEC induction mimics brain development of preterm infants, thus making it a valid model for understanding neurodevelopmental effects of NEC [16].

### Summary

In conclusion, our titratable neonatal mouse model of NEC enables researchers to study a wide range of NEC stages, including mild NEC. This model will facilitate the identification of pathways involved in the progression of NEC, systemic inflammation, and sepsis, as well as cerebral inflammation and disruption of normal neurodevelopment. These data may identify targets for preventive and therapeutic strategies and drug targets.

## Abbreviations

NEC: Necrotizing enterocolitis
DSS: dextran sodium sulfate
GI: gastrointestinal

## Acknowledgements

The authors would like to thank Drs. Gail Besner, Amalia Napoli, Vincent Yang, Ashwini Phadnis-Moghe, Miguel Maderia, and Yijie Wang for scientific discussion and assistance with this manuscript. The authors would also like to thank Amrendra Singh, Noah Teaney, Sarah Bucher, and Stella Ku for their contributions in data collection.

## Data

The authors confirm that the data are availability within the article and its supplementary materials. Additional raw data will be made available upon request.

## Ethical Approval and Consent to participate

Not applicable (no clinical data). All animal experiments were performed with a protocol approved by Stony Brook University IACUC

