## Supplemental Information for "A graded neonatal mouse model of necrotizing enterocolitis demonstrates that mild enterocolitis is sufficient to activate microglia and increase cerebral cytokine expression"

Sha, et al.

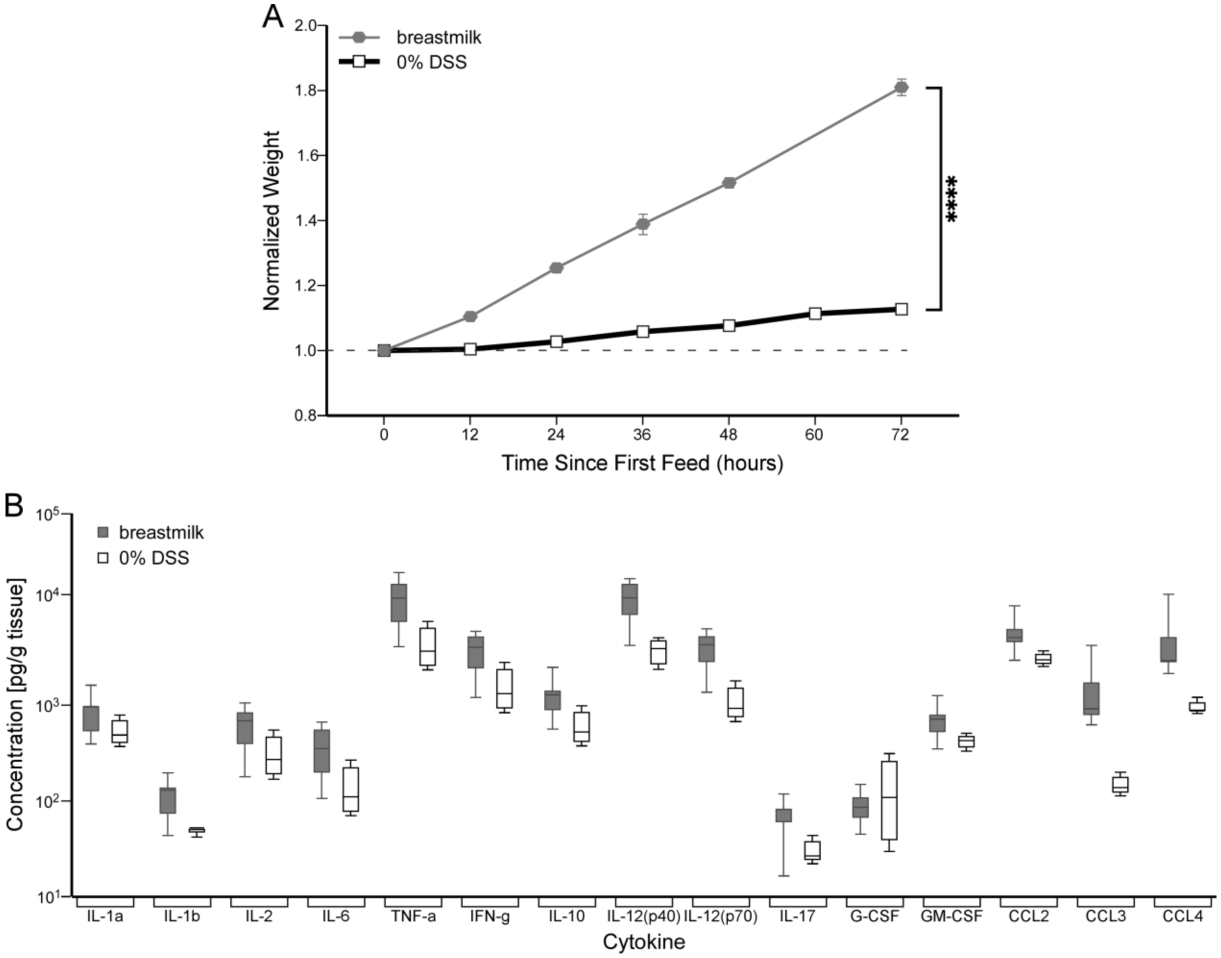

**Supplemental Figure 1. Breastfed mice exhibit notable differences in weight gain and cytokine profile compared to mice fed formula only (0% DSS).**
Breastfed mice gained weight more rapidly compared to formula-fed mice. Breastfed mice nearly doubled their weight within 72 hours. Two-way ANOVA with Tukey's post-hoc. Data presented as mean ± standard error of mean (SEM). ****$p < 0.0001$. Number of mice: breastfed, 7; 0% DSS, 40.

Because of these differences between breastfed and 0% DSS-fed mice, as well as previous findings demonstrating that maternal separation in mice during the first week of life negatively affects neurodevelopment [1], we determined that isolating the mice during feeding presented a confounding variable. Therefore, we used 0% DSS fed mice as a control for our experiments using different supplementations of DSS.

**Supplemental Table 1. Comparisons of Kaplan-Meier survival curves (relates to Figure 1A).**

| | 0% DSS | 0.25% DSS | 1% DSS | 2% DSS | 3% DSS |
|---|---|---|---|---|---|
| 0% DSS | | | | | |
| 0.25% DSS | *0.30* | | | | |
| 1% DSS | *0.73* | *0.23* | | | |
| 2% DSS | ***<0.0001*** | ***0.0054*** | ***<0.0001*** | | |
| 3% DSS | ***<0.0001*** | ***0.0011*** | ***<0.0001*** | ***0.0088*** | |

*P-values* for the comparison between the survival curves of two experimental groups (indicated in the first row and column). A Kaplan-Meier survival analysis was used for mortality during feeding. Significant *p-values* (< 0.05) are in ***bold***.

**Supplemental Table 2. Normalized weights of mice that survived to the final 72-hour timepoint (relates to Figure 1B).**

| | Normalized Weight | | |
|---|---|---|---|
| **Experimental Group** | Mean | SEM | **N (mice)** |
| 0% DSS | 1.13 | 0.01 | 20 |
| 0.25% DSS | 1.14 | 0.02 | 11 |
| 1% DSS | 1.11 | 0.02 | 13 |

Mean weights, standard error of mean (SEM), and counts of all mice that survived the entire feeding protocol.

**Supplemental Table 3. Comparisons of normalized weight curves during feeding (relates to Figure 1B).**

| | 0% DSS | 0.25% DSS | 1% DSS | 2% DSS | 3% DSS |
|---|---|---|---|---|---|
| 0% DSS | | | | | |
| 0.25% DSS | *0.18* | | | | |
| 1% DSS | *0.47* | *0.98* | | | |
| 2% DSS | *0.56* | ***0.031*** | *0.087* | | |
| 3% DSS | ***0.043*** | ***0.0020*** | ***0.0052*** | *0.48* | |

*P-values* for the comparison between weight gain curves of two experimental groups (indicated in the first row and column). A two-way analysis of variance (ANOVA) with Tukey's post-hoc was used for statistical analysis of weight gain curves. Significant *p-values* (< 0.05) are in ***bold***.

**Supplemental Table 4. Clinical Sickness Scores (CSS) of mice during feeding (relates to Figure 1C).**

| | Clinical Sickness Score | | | | |
|---|---|---|---|---|---|
| **Exp Group** | 12 | 24 | 36 | 48 | 60 |
| 0% DSS | 0 ± 0 | 0 ± 0 | 0.07 ± 0.05 | 1.48 ± 0.26 | 1.45 ± 0.28 |
| N (mice) | 29 | 29 | 29 | 23 | 20 |
| 0.25% DSS | 0 ± 0 | 0 ± 0 | 0.05 ± 0.05 | 1.05 ± 0.22 | 0.62 ± 0.31 |
| N (mice) | 24 | 24 | 21 | 19 | 13 |
| 1% DSS | 0 ± 0 | 0 ± 0 | 0.38 ± 0.11 | 1.55 ± 0.27 | 1.87 ± 0.27 |
| N (mice) | 23 | 22 | 21 | 20 | 15 |
| 2% DSS | 0.29 ± 0.18 | 0.29 ± 0.18 | 1.40 ± 0.40 | 2.33 ± 0.33 | 3.00 ± 1.00 |
| N (mice) | 7 | 7 | 5 | 3 | 2 |

Mean ± SEM for the behavior score at each treatment time (in hours) during feeding; the values in the row below are the number of mice alive during that treatment time for behavioral assessment. CSS measures were determined using a scoring system from Zani et al. (2008) [2].

**Supplemental Table 5. Comparisons of CSS during feeding (relates to Figure 1C).**

| | Treatment Time (hours) | | | | | |
|---|---|---|---|---|---|---|
| **Comparison** | 12 | 24 | 36 | 48 | 60 | **Overall** |
| 0% vs 0.25% DSS | *>0.99* | *>0.99* | *>0.99* | *0.18* | ***0.0034*** | ***0.028*** |
| 0% vs 1% DSS | *>0.99* | *>0.99* | *0.38* | *0.99* | *0.27* | *0.29* |
| 0% vs 2% DSS | *0.75* | *0.75* | ***0.0004*** | *0.17* | ***0.012*** | ***<0.0001*** |
| 0.25% vs 1% DSS | *>0.99* | *>0.99* | *0.38* | *0.10* | ***<0.0001*** | ***0.0001*** |
| 0.25% vs 2% DSS | *0.76* | *0.76* | ***0.0004*** | ***0.013*** | ***<0.0001*** | ***<0.0001*** |
| 1% vs 2% DSS | *0.76* | *0.76* | ***0.014*** | *0.24* | *0.12* | ***0.0003*** |

*P-values* for the comparison of behavioral score between two groups at each treatment time (in hours). A two-way ANOVA with Tukey's post-hoc test was used for statistical analysis of the behavioral scores during feeding. Significant *p-values* (< 0.05) are in ***bold***.

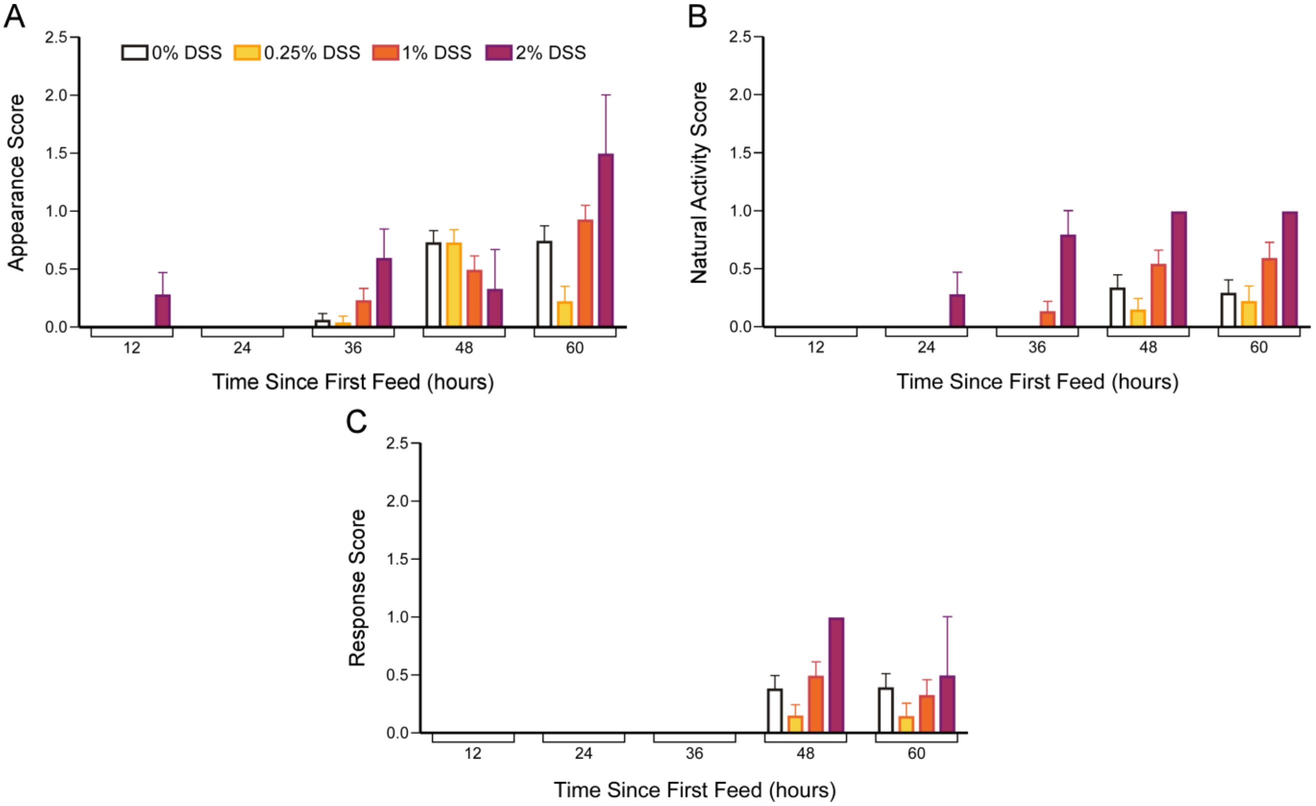

**Supplemental Figure 2. CSS, in each category, show the same overall trend during feeding as the combined CSS (relates to Figure 1C).**

CSS measures were determined using a scoring system from Zani et al. (2008) [2]. Data presented for each score component and stratified by feeding condition. **(A)** Appearance, **(B)** Natural activity, and **(C)** Response to touch scores increase over the course of the feeding protocol, even when animals are fed formula. Furthermore, higher concentrations of DSS are associated with worse CSS earlier. Two-way ANOVA with Tukey's post-hoc, $p < 0.0001$ for all analyses. Data presented as mean ± SEM. Number of mice: 0%, 29; 0.25%, 26; 1%, 26; 2%, 7.

**Supplemental Table 6. External bowel scores of control and DSS-fed mice (relates to Figure 2B).**

| | External Bowel Score | | |
|---|---|---|---|
| **Experimental Group** | Mean | Standard Error of Mean | **N (intestines)** |
| 0% DSS | 0.6 | 0.2 | 18 |
| 0.25% DSS | 1.2 | 0.2 | 13 |
| 1% DSS | 4.0 | 0.3 | 18 |
| 2% DSS | 4.7 | 0.4 | 7 |

Mean external bowel scores, SEM, and counts of mice whose intestines were evaluated macroscopically. Bowels were evaluated using a scoring system from Zani et al. (2008) [2].

**Supplemental Table 7. Comparisons of external bowel scores. (relates to Figure 2B).**

| | 0% DSS | 0.25% DSS | 1% DSS | 2% DSS |
|---|---|---|---|---|
| 0% DSS | | | | |
| 0.25% DSS | *0.33* | | | |
| 1% DSS | ***<0.0001*** | ***<0.0001*** | | |
| 2% DSS | ***<0.0001*** | ***<0.0001*** | *0.27* | |

*P-values* for the comparison of external bowel score between two groups (indicated in the first row and column). A one-way ANOVA with Tukey's post-hoc test was used for statistical analysis of the external bowel scores. Significant *p-values* (< 0.05) are emphasized in bold.

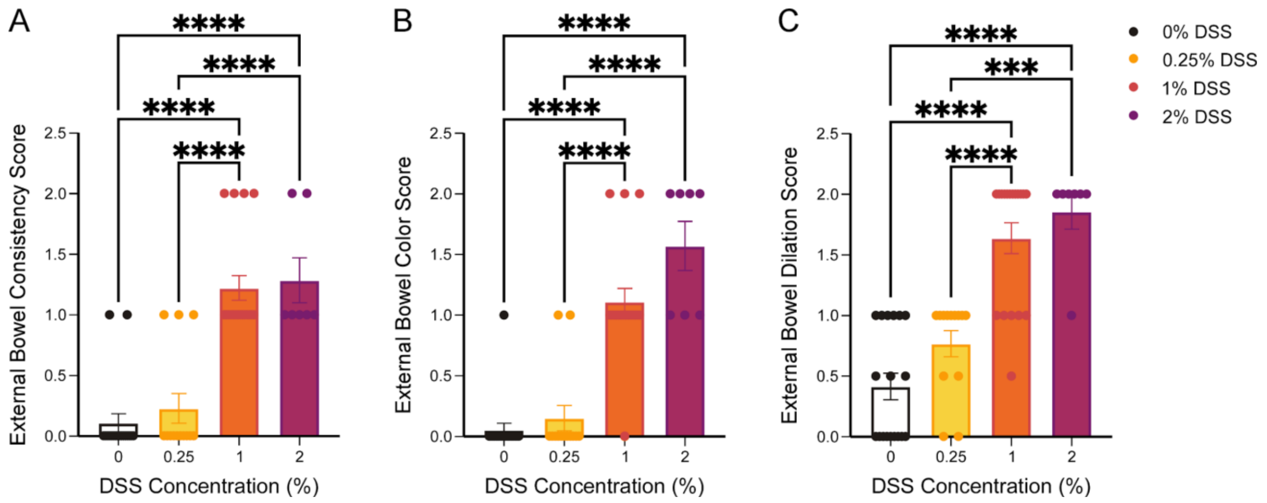

**Supplemental Figure 3. External bowel scores, by category, show the same overall trend as the combined external bowel score (relates to Figure 2B).**
Intestines were assessed as described in Zani et al (2008) [2]. Bowel **(A)** consistency, **(B)** color, and **(C)** dilation are significantly more severe in mice fed with increasing concentrations of DSS (1% and 2% DSS), compared to lower concentrations of DSS (0% and 0.25% DSS). One-way ANOVA with Tukey's post-hoc, $p < 0.0001$ for all analyses. Data presented as mean ± SEM. ***$p < 0.001$, ****$p < 0.0001$. Number of mice: 0%, 18; 0.25%, 13; 1%, 18; 2%, 7.

**Supplemental Table 8. Comparisons of intestinal pathology scores (relates to Figure 2C).**

| | Region of Gastrointestinal (GI) Tract | | |
|---|---|---|---|
| **Comparison** | Proximal Small Bowel | Colon | **Overall** |
| 0% vs 0.25% DSS | ***<0.0001*** | *0.91* | ***0.0005*** |
| 0% vs 1% DSS | ***0.016*** | *>0.99* | *0.18* |
| 0% vs 2% DSS | ***0.0009*** | *0.95* | ***0.0062*** |
| 0.25% vs 1% DSS | *0.88* | *0.95* | *0.70* |
| 0.25% vs 2% DSS | *>0.99* | *>0.99* | *>0.99* |
| 1% vs 2% DSS | *0.96* | *0.96* | *0.84* |

*P-values* for the comparison of intestinal pathology score between two groups at each region of the GI tract. A two-way ANOVA with Tukey's post-hoc test was used for statistical analysis of the intestinal pathology scores. Significant *p-values* (< 0.05) are in ***bold***.

**Supplemental Table 9. Ki-67+ cell counts per small intestinal crypt in mice (relates to Figure 2D).**

| | Ki-67+ Cell Counts per Crypt | | |
|---|---|---|---|
| **Experimental Group** | Mean | SEM | **N (crypts)** |
| 0% DSS | 13.2 | 0.9 | 27 |
| 0.25% DSS | 14.7 | 0.8 | 42 |
| 1% DSS | 13.2 | 1.0 | 16 |
| 2% DSS | 10.9 | 1.1 | 7 |

Mean Ki-67+ cell count, SEM, and counts of all intestinal crypts counted.

**Supplemental Table 10. Comparisons of Ki-67+ scores (relates to Figure 2D).**

| | 0% DSS | 0.25% DSS | 1% DSS | 2% DSS |
|---|---|---|---|---|
| 0% DSS | | | | |
| 0.25% DSS | *0.64* | | | |
| 1% DSS | *0.96* | *0.40* | | |
| 2% DSS | ***0.019*** | ***0.0008*** | *0.091* | |

*P-values* for the comparison of Ki-67+ small intestinal crypt counts between two groups (indicated in the first row and column). A one-way ANOVA with Tukey's post-hoc test was used for statistical analysis of the Ki-67+ counts. Significant *p-values* (< 0.05) are in ***bold***.

**Supplemental Table 11. Linear regression with log-transformation analysis of the concentrations of cytokines and chemokines in liver tissue (relates to Figures 3 and 4).**

| Cytokine | Slope (m) | Y-intercept (b) | R-squared | F-statistic | P-value |
|---|---|---|---|---|---|
| IL-1α | 0.21 | 1.3 | 0.30 | 21 | ***<0.0001*** |
| IL-1β | 0.098 | 0.61 | 0.091 | 4.8 | ***0.033*** |
| IL-2 | 0.20 | 1.2 | 0.33 | 24 | ***<0.0001*** |
| IL-6 | 0.18 | 0.99 | 0.21 | 13 | ***0.0009*** |
| IL-10 | 0.24 | 1.5 | 0.36 | 26 | ***<0.0001*** |
| IL-12(p40) | 0.098 | 2.9 | 0.11 | 5.8 | ***0.020*** |
| IL-12(p70) | 0.31 | 2.5 | 0.41 | 33 | ***<0.0001*** |
| IL-17 | 0.15 | 0.33 | 0.16 | 9.4 | ***0.0035*** |
| G-CSF | 0.63 | 0.73 | 0.42 | 34 | ***<0.0001*** |
| GM-CSF | 0.95 | 0.97 | 0.18 | 10 | ***0.0024*** |
| IFN-γ | 0.24 | 1.9 | 0.35 | 26 | ***<0.0001*** |
| CXCL1 | 0.13 | 1.7 | 0.093 | 4.9 | ***0.031*** |
| CCL2 | 0.11 | 2.2 | 0.098 | 5.2 | ***0.027*** |
| CCL3 | 0.35 | 0.47 | 0.26 | 17 | ***0.0002*** |
| CCL4 | 0.23 | 1.6 | 0.37 | 28 | ***<0.0001*** |
| TNF-α | 0.22 | 2.3 | 0.33 | 23 | ***<0.0001*** |

Liver cytokine and chemokine concentrations were first log-transformed using the equation y = log(y) and then analyzed using simple linear regression. The results of the linear regression analysis are summarized in the table and graphically displayed in Figures 3 and 4. The slopes of the linear regression lines were statistically compared to the line y = 0, using a two-tailed t-test, to determine if there is a significant correlation. Significant *p-values* (< 0.05) are in ***bold***.

**Supplemental Table 12. Linear regression with log-transformation analysis of the concentrations of cytokines and chemokines in blood plasma (relates to Figure 5).**

| Cytokine | Slope (m) | Y-intercept (b) | R-squared | F-statistic | *p*-value |
|---|---|---|---|---|---|
| IL-1α | -0.040 | 1.4 | 0.0011 | 0.046 | *0.83* |
| IL-1β | *below detection threshold* | | | | |
| IL-2 | *below detection threshold* | | | | |
| IL-6 | 0.45 | 0.22 | 0.078 | 3.5 | *0.070* |
| IL-10 | 0.45 | 0.21 | 0.34 | 21 | ***<0.0001*** |
| IL-12(p40) | 0.27 | 3.2 | 0.17 | 8.2 | ***0.0066*** |
| IL-12(p70) | *below detection threshold* | | | | |
| IL-17 | *below detection threshold* | | | | |
| G-CSF | 0.64 | 2.2 | 0.22 | 12 | ***0.0016*** |
| GM-CSF | *below detection threshold* | | | | |
| IFN-γ | *below detection threshold* | | | | |
| CXCL1 | 0.63 | 0.84 | 0.22 | 11 | ***0.0016*** |
| CCL2 | *below detection threshold* | | | | |
| CCL3 | 0.57 | 0.10 | 0.088 | 3.9 | *0.054* |
| CCL4 | 0.63 | 1.2 | 0.18 | 9.1 | ***0.0045*** |
| TNF-α | *below detection threshold* | | | | |

Same as in Supplemental Table 11 but for plasma cytokines and chemokines. The results of the simple linear regression analysis are graphically displayed in Figure 5 and Supplemental Figure 4.

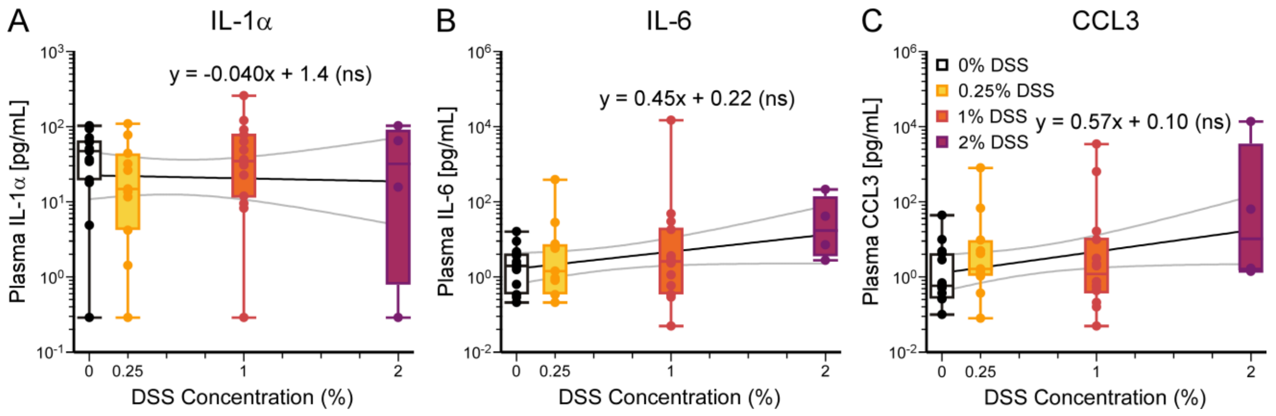

**Supplemental Figure 4. Certain blood plasma cytokine and chemokine concentrations do not significantly correlate with DSS concentration (relates to Figure 5).**

Cytokines include **(A)** IL-1α, $p = 0.83$, **(B)** IL-6, $p = 0.070$, and **(C)** CCL3, $p = 0.054$. All other cytokines (IL-1β, IL-2, IL-12(p70), IL-17, GM-CSF, CCL2, IFN-γ, TNF-α) tested in plasma were below the threshold for detection. Simple linear regression with log-transformation of y values performed (see Supplemental Table 12). Data presented as boxplots showing min-max. Slope and y intercept with confidence intervals are plotted. *ns* = not significant ($p \geq 0.05$). Number of mice: 0%, 14; 0.25%, 11; 1%, 14; 2%, 4.

**Supplemental Table 13. Linear regression with log-transformation analysis of the concentrations of cytokines and chemokines in brain tissue (relates to Figure 6).**

| **Cytokine** | **Slope (m)** | **Y-intercept (b)** | **R-squared** | **F-statistic** | ***p*-value** |
|---|---|---|---|---|---|
| IL-1α | 0.17 | 0.32 | 0.11 | 3.5 | *0.074* |
| IL-1β | 0.084 | -0.30 | 0.046 | 1.4 | *0.25* |
| IL-2 | 0.18 | 0.014 | 0.13 | 4.5 | ***0.043*** |
| IL-6 | 0.083 | -0.10 | 0.018 | 0.54 | *0.47* |
| IL-10 | 0.12 | 0.21 | 0.073 | 2.3 | *0.14* |
| IL-12(p40) | 0.0018 | 2.4 | 0.000014 | 0.00042 | *0.98* |
| IL-12(p70) | 0.044 | 1.2 | 0.0039 | 0.11 | *0.75* |
| IL-17 | 0.12 | -0.70 | 0.093 | 3.0 | *0.096* |
| G-CSF | 0.49 | 0.45 | 0.27 | 11 | ***0.0030*** |
| GM-CSF | 0.17 | 0.28 | 0.10 | 3.1 | *0.089* |
| IFN-γ | 0.15 | 0.62 | 0.085 | 2.7 | *0.11* |
| CXCL1 | 0.21 | 1.2 | 0.27 | 10 | ***0.0035*** |
| CCL2 | 0.11 | 1.6 | 0.039 | 1.2 | *0.29* |
| CCL3 | -0.13 | 0.81 | 0.077 | 2.4 | *0.13* |
| CCL4 | 0.091 | 0.78 | 0.019 | 0.57 | *0.46* |
| TNF-α | 0.0059 | 0.97 | 0.00018 | 0.0051 | *0.94* |

Same as in Supplemental Table 11 but for brain cytokines and chemokines. The results of the simple linear regression analysis are graphically displayed in Figure 6 and Supplemental Figures 5 and 6.

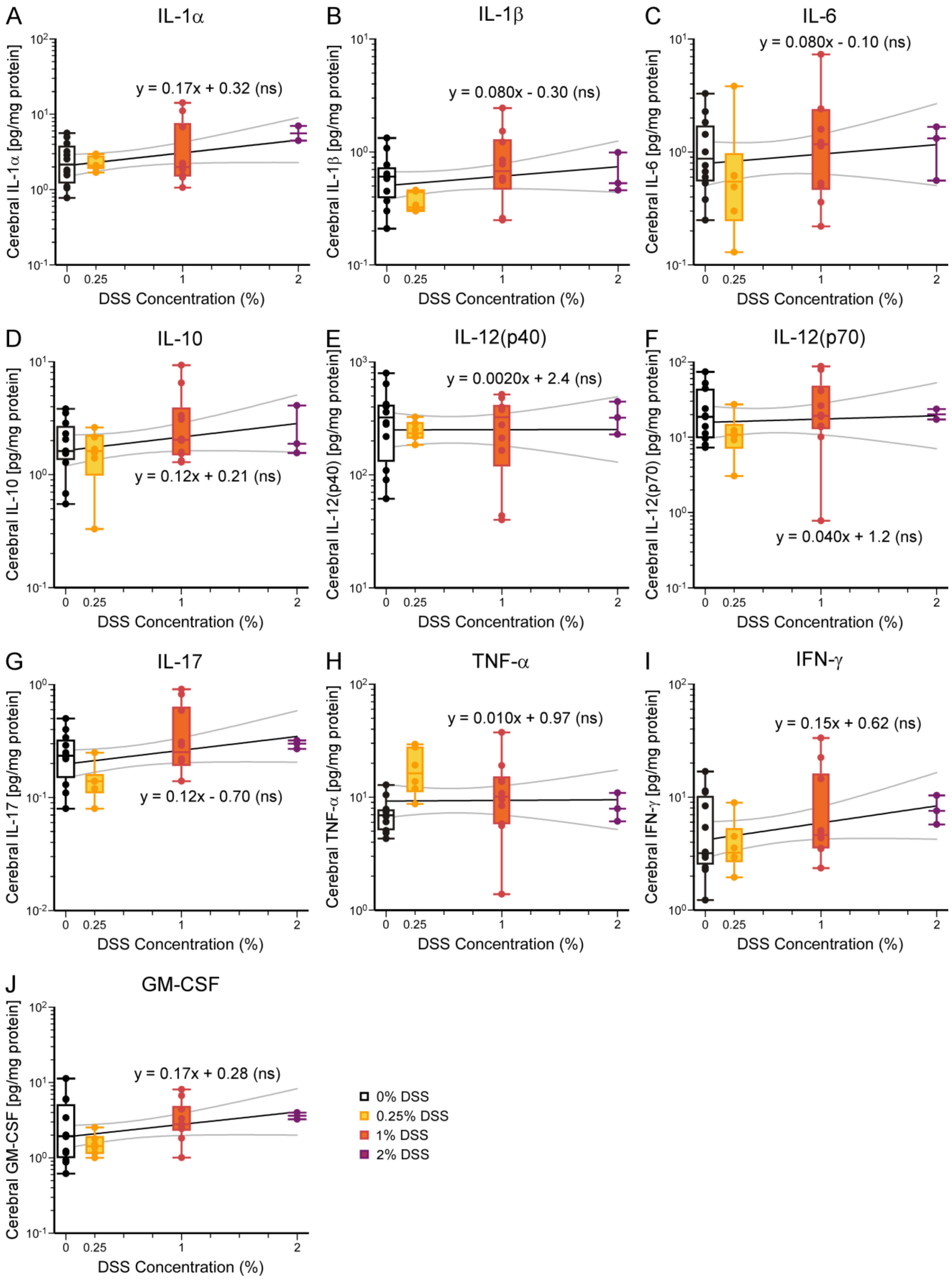

A
IL-1α
y = 0.17x + 0.32 (ns)
Cerebral IL-1α [pg/mg protein]
B
IL-1β
y = 0.080x - 0.30 (ns)
Cerebral IL-1β [pg/mg protein]
C
IL-6
y = 0.080x - 0.10 (ns)
Cerebral IL-6 [pg/mg protein]
D
IL-10
y = 0.12x + 0.21 (ns)
Cerebral IL-10 [pg/mg protein]
E
IL-12(p40)
y = 0.0020x + 2.4 (ns)
Cerebral IL-12(p40) [pg/mg protein]
F
IL-12(p70)
y = 0.040x + 1.2 (ns)
Cerebral IL-12(p70) [pg/mg protein]
G
IL-17
y = 0.12x - 0.70 (ns)
Cerebral IL-17 [pg/mg protein]
H
TNF-α
y = 0.010x + 0.97 (ns)
Cerebral TNF-α [pg/mg protein]
I
IFN-γ
y = 0.15x + 0.62 (ns)
Cerebral IFN-γ [pg/mg protein]
J
GM-CSF
y = 0.17x + 0.28 (ns)
Cerebral GM-CSF [pg/mg protein]
DSS Concentration (%)
0% DSS
0.25% DSS
1% DSS
2% DSS

**Supplemental Figure 5. Many brain cytokine concentrations do not significantly correlate with DSS concentration (relates to Figure 6).**

In the brain, concentrations of the cytokines IL-2, $p = 0.043$ (Figure 6A), and G-CSF, $p = 0.003$ (Figure 6B), are positively correlated with DSS concentration. In contrast, other cytokines tested in the brain, did not show the same trend. These cytokines include **(A)** IL-1α, $p = 0.074$, **(B)** IL-1β, $p = 0.25$, **(C)** IL-6, $p = 0.47$, **(D)** IL-10, $p = 0.14$, **(E)** IL-12(p40), $p = 0.98$, **(F)** IL-12(p70), $p = 0.75$, **(G)** IL-17, $p = 0.096$, **(H)** TNF-α, $p = 0.94$, **(I)** IFN-γ, $p = 0.11$, and **(J)** GM-CSF, $p = 0.089$ (see Supplemental Table 13). Results displayed and analyzed as in Supplemental Figure 4. *ns* = not significant ($p \geq 0.05$). Number of mice: 0%, 12; 0.25%, 6; 1%, 10; 2%, 3.

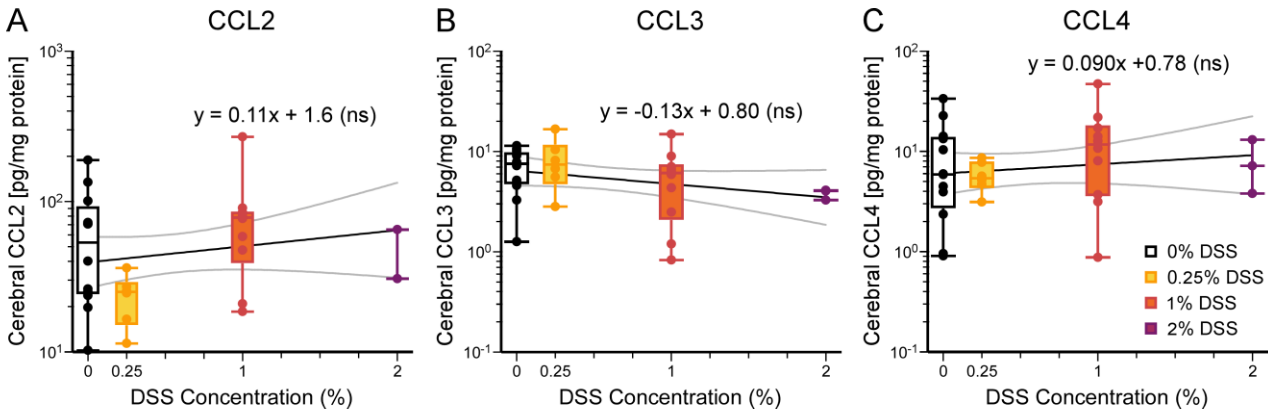

**Supplemental Figure 6. Many Brain chemokine concentrations do not significantly correlate with DSS concentration (relates to Figure 6).**
In the brain, only the concentration of chemokine CXCL1, $p$ = 0.0035 (Figure 6C), is positively correlated with DSS concentration. Other chemokines do not show significant trends in the brain; these chemokines include **(A)** CCL2, $p$ = 0.29, **(B)** CCL3, $p$ = 0.13, and **(C)** CCL4, $p$ = 0.46 (see Supplemental Table 13). Results displayed and analyzed as in Supplemental Figure 4. *ns* = not significant ($p \geq 0.05$). Number of mice: 0%, 12' 0.25%, 6; 1%, 10; 2%, 3.

**Supplemental Table 14. Comparisons of neuron proportions in CA1 hippocampus (relates to Figure 7B).**

| | 0% DSS | 0.25% DSS | 1% DSS | 2% DSS |
|---|---|---|---|---|
| 0% DSS | | | | |
| 0.25% DSS | *>0.99* | | | |
| 1% DSS | *0.49* | *0.63* | | |
| 2% DSS | *0.43* | *0.31* | ***0.027*** | |

*P-values* for the comparison, between two groups (indicated in the first row and column), of the proportion of neurons among all cells in the CA1 hippocampal region. A one-way ANOVA with Tukey's post-hoc test was used for statistical analysis of the neuron proportions. Significant *p-values* (< 0.05) are in ***bold***.

**Supplemental Table 15. Comparisons of microglia proportions in CA1 hippocampus (relates to Figure 7C).**

| | 0% DSS | 0.25% DSS | 1% DSS | 2% DSS |
|---|---|---|---|---|
| 0% DSS | | | | |
| 0.25% DSS | *0.50* | | | |
| 1% DSS | *0.33* | ***0.018*** | | |
| 2% DSS | *0.66* | *0.068* | *0.94* | |

*P-values* for the comparison, between two groups (indicated in the first row and column), of the proportion of microglia among all cells in the CA1 hippocampal region. A one-way ANOVA with Tukey's post-hoc test was used for statistical analysis of the microglia proportions. Significant *p-values* (< 0.05) are in ***bold***.

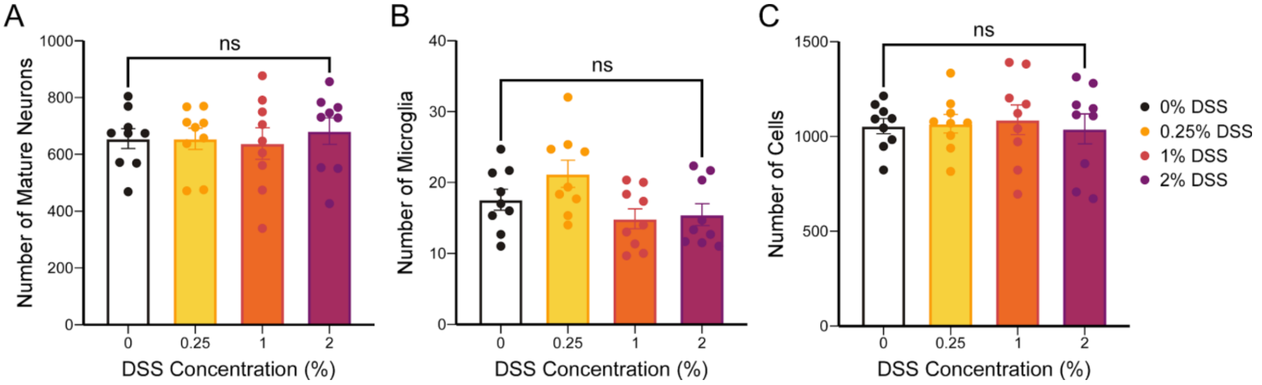

**Supplemental Figure 7. No significant difference in total number of neurons, microglia, or DAPI-positive cells in CA1 hippocampal region seen across all experimental groups (relates to Figure 7).**
**(A)** NeuN-positive cell counts across all DSS groups, $p = 0.3649$.
**(B)** Iba1-positive cell counts across all DSS groups, $p = 0.0694$.
**(C)** DAPI-positive cell counts across all DSS groups, $p = 0.9157$.
One-way ANOVA with Tukey's post-hoc. Data presented as mean ± SEM. *ns* = not significant ($p \geq 0.05$). n = 9 immunohistochemical images for all experimental groups.
